# Divergence of root system traits in soybean between breeding and diversity lines

**DOI:** 10.1101/2025.05.05.651701

**Authors:** Sujata Bogati, Joshua Carpenter, Jinha Jung, Sam Schafer, Jairam Danao, Ellen Woods, Qijian Song, Michael Kantar, Jianxin Ma, Diane R. Wang

## Abstract

Roots are critical for supporting basic plant functions such as anchoring in various substrates, uptake of water and nutrients, and hosting symbiotic relationships. In crops, indirect changes to root system architecture (RSA) have occurred largely as a result of selection for yield or other related aboveground traits. In cultivated soybean (*Glycine max*), evidence of changes to RSA resulting from breeding for crop performance has been inconsistent, with some studies supporting an overall decrease in performance-related trait values, such as root length and density, and other work showing the opposite. The current study sets out to ask whether there is any systematic differentiation in RSA between a set of elite breeding lines (n=8) of soybean developed for the Midwest United States and a group of biogeographically diverse landraces from the USDA Soybean Germplasm Collection (n=16. Groups are compared across three distinct developmental stages (V2–V6, V7–R2, R3–R7) and two contrasting soil environments. In total, 432 root systems were phenotyped for 12 structural traits derived from 2D images along with root and shoot biomass. A new 3D root modeling approach leveraging photogrammetry-derived pointclouds is additionally tested on a subset of 30 contrasting root systems. Results indicate that the diversity lines had smaller root systems overall but greater phenotypic plasticity in response to soil environment as compared to breeding lines. Additionally, the study finds evidence for trade-offs between above-ground and below-ground trait plasticity.

**Core ideas:**

- Previous work suggests crop root system architecture (RSA) has changed as a result of indirect selection
- This study compares RSA in elite breeding lines with that of diverse landraces in cultivated soybean
- A total of 432 root systems are analyzed across 24 genotypes, two soil environments and three developmental stages
- Findings show diversity lines have smaller root systems but greater plasticity in response to soil environment

## Introduction

The root system provides essential functions for the whole plant. These include anchoring, facilitating symbiotic partnerships with soil microorganisms, and serving as the initial conduit for nutrient and water uptake (McNear, 2013). Understanding the relationships between root traits and their connections to above-ground features has been a key area of interest in the plant sciences (O’Toole & Bland, 1987; Watt et al. 2006). Recently, there has been growing focus on harnessing root traits to enhance crop resource utilization (Lynch, 2019, Tracy et al., 2020); this is especially critical as climate change negatively impacts soil conditions, motivating the development of more resilient crop varieties that can thrive with fewer inputs.

Modifications to the root system as part of varietal crop improvement have largely occurred through indirect selection on yield and other above-ground performance metrics rather than direct selection on root traits themselves (York et al., 2015). For instance, in commercial maize hybrids, empirical breeding for yield has resulted in smaller root systems over the course of 80 years (Rinehart *et al.*, 2024). Similarly, modern high-yielding wheat varieties boast fewer roots per plant than historic varieties (Fradgley et al., 2020). In rice, deeper rooting, conferred by higher expression of the *DRO1* gene, gives rise to greater yields under drought conditions (Uga *et al.*, 2013). These kinds of insights highlight the important role that root systems can play in crop improvement, especially for resource-limited environments, and strategies to incorporate direct selection on root traits by combining laboratory and field phenotyping have been proposed (Wasson et al., 2012). However, several challenges to successful direct selection of root system traits by breeding programs remain, including the relatively low heritabilities of root-related traits that are associated with strong genotype-by-environment (G x E) interactions and phenotypic plasticity (Robinson, 2001; Tuberosa et al., 2002; Malamy, 2005; Comas et al., 2013).

Genotype-by-environment interactions refer to the differential responses of genotypes to varied environments. These interactions can complicate varietal selection, especially if they result in genotype rank changes across environment (e.g., locations or years). G x E is conceptually and functionally related to phenotypic plasticity, which refers to modification of traits generally of single genotypes in response to change(s) to the environment. Plasticity is conferred by changes in physiology, anatomy, morphology, and/or resource allocation and is regarded as genetically controlled (Bradshaw, 2006; Sultan, 2000). In the context of varietal crop improvement, greater phenotypic plasticity has been considered by some to be promising for low-input systems, enabling genotypes to acclimate to stress scenarios (Lynch, 2019; Schneider and Lynch, 2020). However, it is important to note that plasticity at one level of biological organization may be responsible for stability at another. Moreover, greater plasticity is not inherently adaptive and may simply the inevitable consequences of resource limitation (Van Kleunen and Fischer 2005; Ghalambor et al. 2007;Sultan, 2015).

In soybean (*Glycine max* L. Merr.), a globally important source of plant-based protein for both humans and animals, there is conflicting evidence for the effects that modern breeding has had on root systems. Recent work comparing 11 genotypes, including elite breeding lines and exotic germplasm, for root traits under rainfed conditions suggested that a parsimonious root system, characterized by reduced number of axial roots and reduced lateral root length and density, is beneficial under high-input environments and a potential reason for superior biomass formation in elite breeding lines (Noh et al., 2022). These results contrast with those from another study, which directly examined root system traits across soybean varieties released across different eras; it suggested that root biomass, root volume, root length, root surface area, and root-shoot ratio increased over time (Mandozai et al., 2021). One complicating factor for studies on soybean root variation is that root development is linked to plant growth stages, with distinct patterns emerging throughout the plant’s lifecycle. During vegetative growth, rapid root growth occurs, and as the plant transitions to early reproductive stages, root branching becomes more prominent. Finally, root growth generally decreases during pod development (Hoogenboom et al., 1987). Despite the impact of overall plant development on root traits, research that explicitly considers developmental stages has been limited, and more comprehensive studies that sample roots across multiple growth stages are needed (Ordonez et al., 2020).

Developments in phenotyping methods that reduce the labor involved in collecting root trait measurements open up new opportunities for addressing fundamental questions about how root trait variation is partitioned across genetics, environments, and growth stages. Recent advancements have enhanced researchers’ ability to understand key root traits that benefit crops, aiding in the selection of improved varieties (Kuijken et al. 2015; Song et al. 2021). 2D imaging using digital cameras is now considered a conventional approach in root phenotyping, with software platforms readily available to researchers for image analysis such as WinRhizo (2016a, Regent Instruments, Quebec, Canada) and open-source ImageJ (Schneider et al. 2012). The integration of modern imaging technologies, such as MRI, X-ray CT, and PET, also enables more precise 3D measurements than was possible before (Atkinson et al., 2019; Messina et al., 2021; Stingaciu et al., 2013). However, portable 3D imaging approaches that do not rely on expensive technologies are not yet widely accessible to researchers, and 2D imaging remains the standard approach.

In the current study, we compared root traits among 24 diversity and breeding lines of soybean across three growth stages at two locations with differing soil environments. In total, 432 individual soybean root systems were collected and 2D imaged to extract root architectural traits. To address the need for more cost-effective 3D phenotyping approaches, we additionally tested the utility of a new analytical framework to reconstruct 3D root models from 2D images collected economically using a common digital single-lens reflex (DSLR) camera and a portable turn-table phenotyping system. Our overall objectives were to: (1) determine whether there were differences in root traits between breeding lines and diversity lines, (2) quantify the relative effects of genotype group, environment, and their interaction on root traits, and (3) compare features derived from 3D root models with traits measured using conventional 2D image analysis.

## Materials and methods

*Plant materials.* Twenty-four soybean genotypes were evaluated for this study (**Table S1**). Sixteen of these, hereafter referred to as the diversity lines (DL), were selected from the USDA Soybean Germplasm Collection and originated from seven different countries (United States, Japan, China, South Korea, Russia, Vietnam, and the Netherlands) (**Figure 1**).

**Figure 1.**
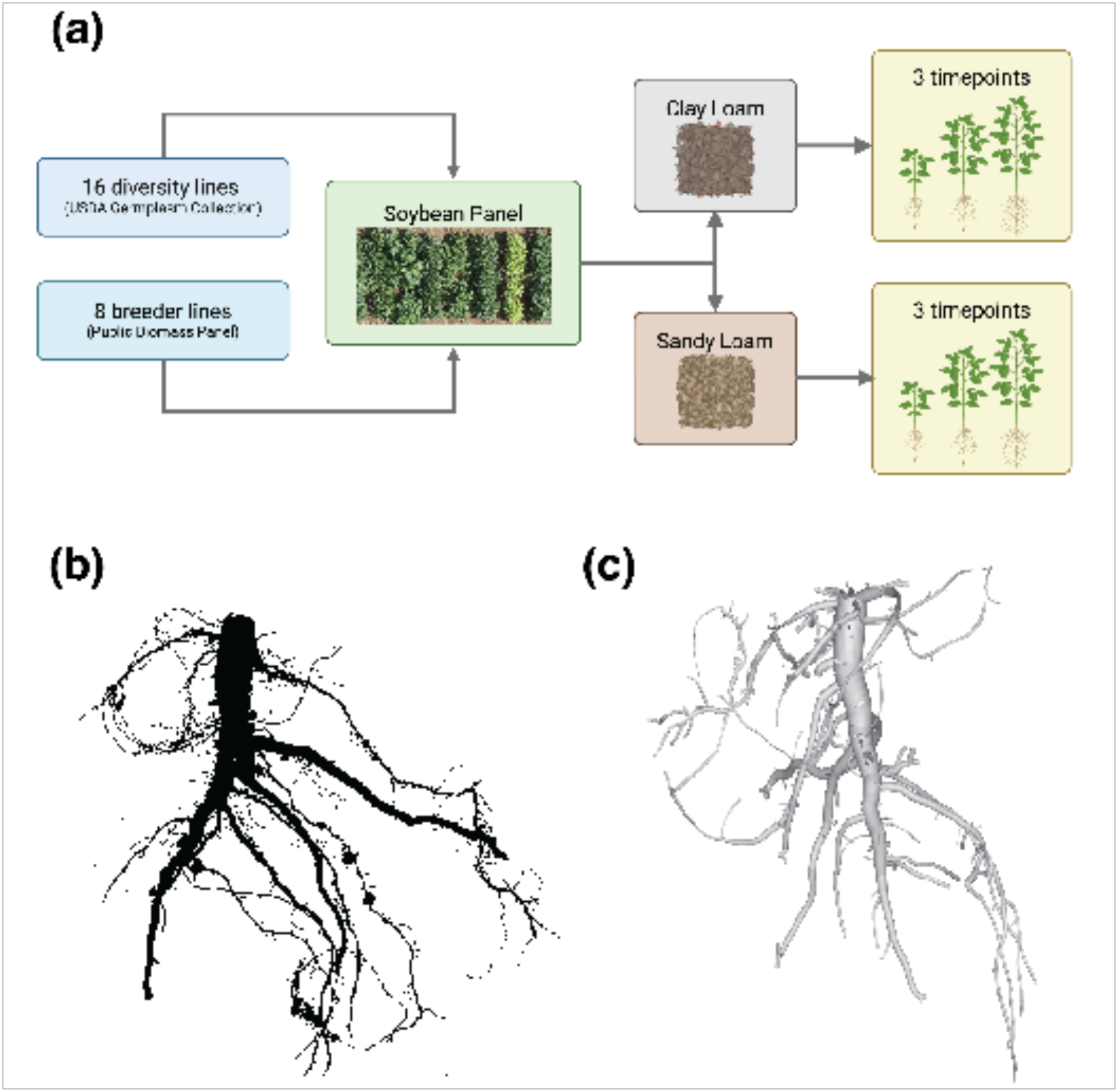
Study overview. **(a)** Diagram showing the study’s experimental design with 24 soybean genotypes classified into two genotype groups (diversity lines and breeder lines), two soil environments (clay loam and sandy loam), and three developmental timepoints. All root samples were phenotyped using 2D imaging (n = 432), and a subset (n = 30) was additionally measuring using a 3D reconstruction approach. **(b)** An example of a 2D root image taken of one of the breeding lines, G5, sampled from timepoint two in the clay loam site. **(c)** An example from 3D reconstruction of the same root sample.

These accessions were selected from a larger set of 250 USDA accessions that were initially evaluated in 2021 at the Pinney Purdue Agricultural Center (Wanatah, Indiana). The 16 DL were able to flower and mature under Indiana conditions and were selected based on phenotypic variation of several above-ground variables, including specific leaf area, plant height and leaf-level stomatal conductance. Eight additional lines, hereafter referred to as the Breeding lines (BL), were additionally selected for this study. The BL originated from a larger breeder-nominated set of germplasm known as the Public Biomass Panel, which was comprised of germplasm developed for production in the Midwest U.S. These additional eight BL were selected to represent the range of phenotypic variation observed in the full panel based on preliminary data gathered in 2021 also for specific leaf area, plant height and leaf-level stomatal conductance.

*Genotyping*: All DLs were previously genotyped using SoySNP50K, a 50K genome-wide single nucleotide polymorphism (SNP) beadchip. The BLs were genotyped for the current study using the BARCSoySNP6K Assay, a 6K genome-wide SNP beadchip (Song et al. 2013, 2015); critically, these 6K SNPs were a direct subset of SoySNP50K, enabling combined analyses with DLs. The GenomeStudio Genotyping Module v2.0.5 (GenomeStudio Software Downloads (illumina.com) was used to extract the genotypes from 6K. A customized R script was used to merge 6k SNPs with 50k dataset based on position as both the files were aligned to soybean reference genome Wm82.a2 (https://data.soybase.org/Glycine/max/diversity/Wm82.gnm1.div.Song_Hyten_2015/glyma.Wm82.gnm1.div.Song_Hyten_2015.vcf.gz).

*Experimental design and data collection.* The study was carried out during the growing season (May through October) of 2022 at the Pinney Purdue Agricultural Center (PPAC) in Wanatah, Indiana (longitude: -86.928948, latitude: 41.442025) located in Indiana’s northwest region (**Figure S1a**). During the growing season, PPAC had an average air temperature of 18°C and an average soil temperature of 19.5 °C (Figure S1b). Cumulative precipitation during the season was approximately 423.7 mm (about 16.68 in). The experiment was conducted at two locations (environments) within PPAC approximately 2.6 km apart. The locations differed in soil properties; one was characterized as a sandy loam while the other was a clay loam. Soil analysis (soil nutrients organic matter, phosphorous, potassium, magnesium, calcium, sodium, copper, zinc, sulfur and boron), soil texture (sand, silt, clay), bulk density, water holding capacity, soil pH, buffer pH, cation exchange capacity) were carried out at the A & L Great Lakes Laboratories (Fort Wayne, Indiana, USA) prior to planting. In addition to soil texture, the two sites differed according to water holding capacity, cation exchange capacity, and several nutrients (Figure S1c).

Within each location, fields were planted following a randomized complete block design with 24 genotype entries and three blocks. Each site was planted in single-row plots that were 3 m in length (2.1 m short rows with 0.9 m alley). Seeds were hand planted to a depth of approximately 2.5 cm because of limited seed availability and the small plot sizes. The previous crop was corn at both sites. Across all genotypes, plants were destructively sampled from both sites at three time points—TP1 (June 27^th^ – July 14^th^), TP2 (July 15^th^ – August 3^rd^), and TP3 (August 4^th^ – August 18^th^)—roughly corresponding to growth stages V2–V6 (second to sixth trifoliate), V7–R2 (seventh trifoliate [corresponding to early flowering in most genotypes] to full bloom), and R3–R7 (beginning pod to beginning maturity), respectively. Each site was planted in single-row plots that were 3 m in length (2.1 m short rows with 0.9 m alley). Seeds were hand planted to a depth of approximately 2.5 cm because of limited seed availability and the small plot sizes. The previous crop was corn at both sites. Prior to any losses, the full experimental sample consisted of 432 plants (24 lines x 3 spatial blocks/site x 2 sites x 3 timepoints). During each sampling event, one plant was selected randomly per plot, avoiding border plants. Individual plants were excavated with shovels, taking care to minimize damage to the root systems as well as those of neighboring experimental plants. Roots and shoots were separated and dried: shoot samples were dried at 60°C and weighed, while root samples were washed with water and then air-dried. Once dried, root samples were stored in the freezer in Ziploc bags. After the field season ended, imaging and trait analysis was performed on clean, dry samples.

*2D imaging and trait extraction.* Images were collected using an imaging platform comprised of a copy stand (Kaiser RS 2 CP Copy Stand), light box, DLSR camera (Canon EOS Rebel T7 DSLR Camera with EF-S 18-55mm f/3.5-5.6 IS II lens; Canon Inc., Japan, Taiwan, and China), and clear plastic trays on which samples were spread (**Figure S2**). Root samples were placed on trays oriented to minimize the occlusion of lateral roots. Resultant digital images were analyzed using both WinRHIZO (2016a, Regent Instruments, Quebec, Canada) and ImageJ version 2.16.0/1.54p (https://imagej.net/ij/). WinRHIZO was used to automatically detect and segment root samples and quantify six traits using built-in analysis options (**Table 1**). Using ImageJ, an additional six traits were quantified (**Table 1**). For tap root diameter, four to five random diameters across the tap roots were measured, and the maximum was used for eventual analysis. Similarly, four to five random root angles were measured in ImageJ and the average root angle was computed for each sample. For lateral branch length, four to five random lateral branches were selected in ImageJ and their lengths were measured; the maximum value was eventually used for subsequent analyses. Additional details on ImageJ trait extraction are provided in **Figure S3**.

**Table 1.**
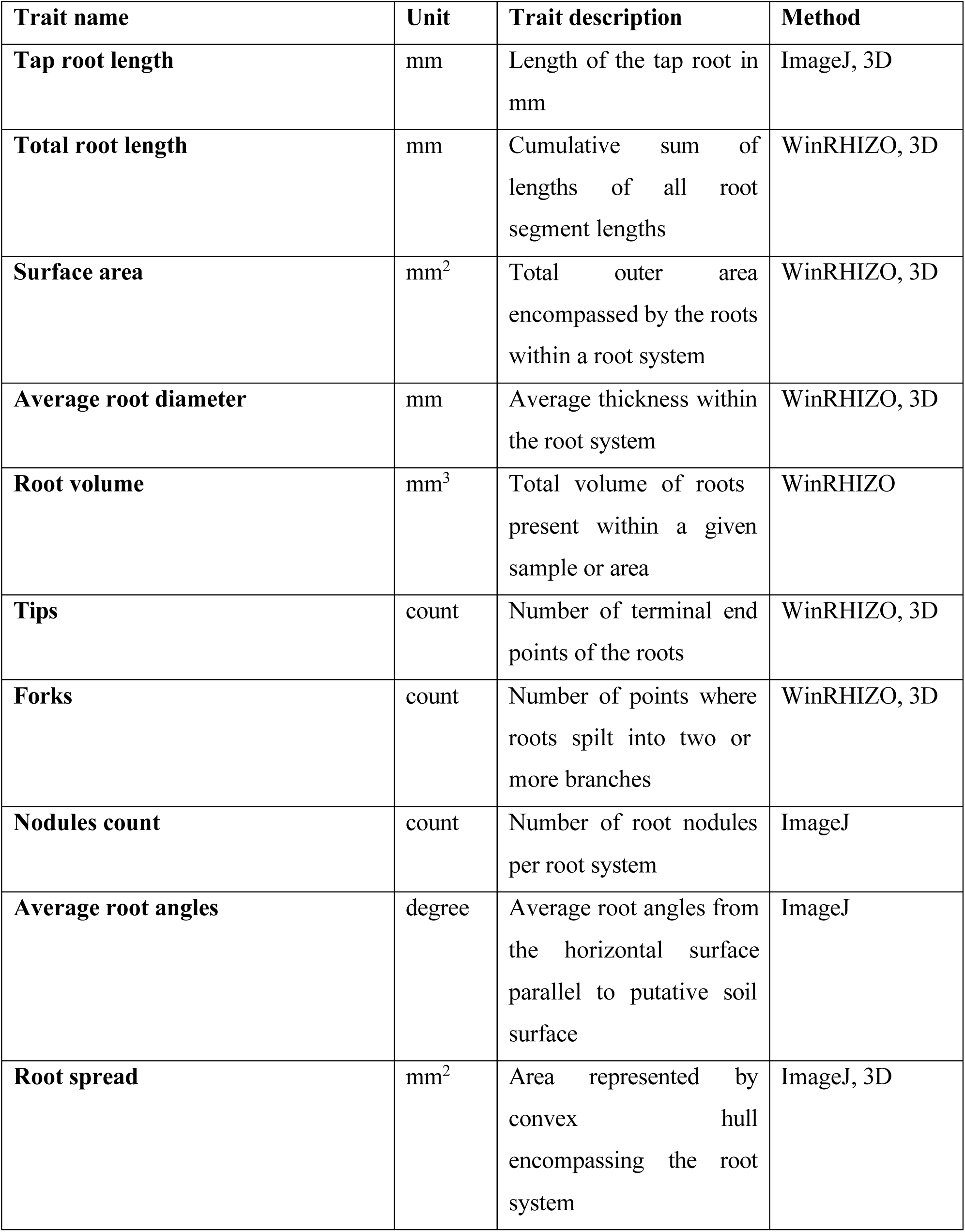
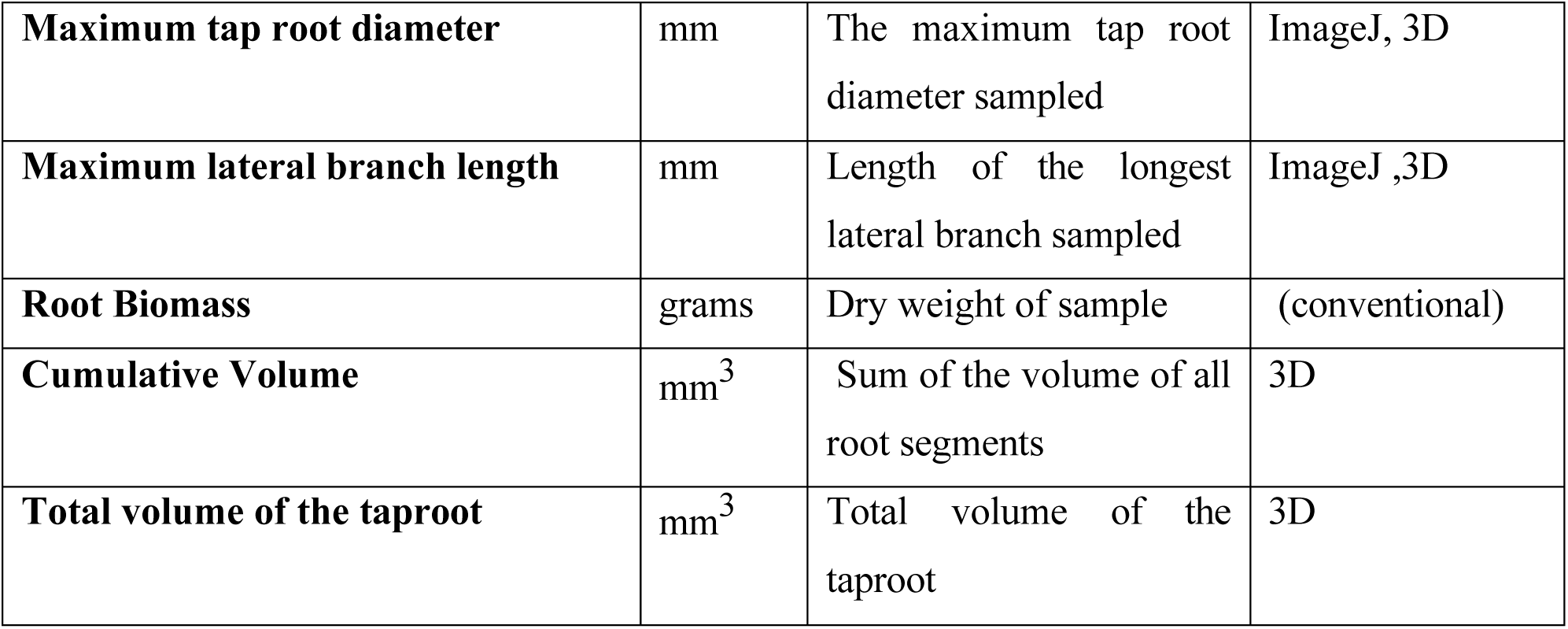
Summary of root traits measured.

*3D imaging and trait extraction:* To assess the feasibility and usefulness of 3D modeling, a subset of the roots was selected for 3D scanning (n=30). From each developmental timepoint, approximately equal samples from each soil environment were selected, randomly sampling across genotypes. Close range photogrammetric scanning was used to generate an initial 3D point cloud of the selected samples. Each root sample was clamped into a holder on a turntable (Figure S2). As the turntable rotated, a Sony DLSR camera (Sony Alpha a7 II with 24.3MP full-frame Exmor CMOS sensor; Sony Corporation, Japan and Thailand) captured photos every 5 degrees around the sample at a resolution of 5304px by 7952px. A small aperture (f/22) was maintained to create the largest depth of field possible during the image capture process. This allowed the lateral branches of each root to remain in focus even as they rotated towards and away from the camera position. After capture, Agisoft’s Metashape was used to implement structure-from-motion (SfM) 3D reconstruction. This method of reconstruction produced a scaleless point cloud of each root sample. The number of points varied widely by the size of the root. Samples from the first timepoint were represented by approximately 200-thousand points, while samples from the last timepoint were represented by nearly 6-million points. We scaled the point cloud to correct dimensions using scale references attached to the turntable prior to imaging. After 3D reconstruction, a hierarchical cylinder model of each root was generated from the point cloud.

### Statistical analysis

Results were visualized using either R version 4.4.1 (R-Core Team, 2023) with the *ggplot* package (Wickham, 2016) or Python version 3.10.12 (Van Rossum and Drake, 2009) with the packages, *Matplotlib* version 3.4.2 (Hunter, 2007) and *seaborn* version 0.11.2 (Waskom, 2011). For SNP data, Principal Components Analysis (PCA) and hierarchical clustering analysis were carried out using *FactoMineR* (Lê *et al.*, 2008) and base R, respectively, with the ‘complete’ agglomeration method, respectively. For trait data, Principal Components Analysis was carried out with Python library *scikit-learn* version 1.3.2 (Pedregosa et al., 2011) using the ‘sklearn.decomposition’ module. Trait data were first averaged for each genotype by timepoint combination, and variables were standardized prior to calculation of the covariance matrix for PCA. Fisher’s Linear Discriminant Analysis (LDA) was carried out to determine the relative contribution of root traits and shoot biomass to the separation of breeder and diversity lines. This process was applied to each timepoint separately. To account for different scales, predictors were standardized prior to applying LDA. Target classes for separation were the two genotype categories, *i.e.*, breeder lines and diversity lines. Five-fold cross-validation was used for model development, and average LDA coefficients across the five folds were calculated for each predictor for each timepoint. These coefficients were visualized as heatmaps to display the relative importance of each trait in distinguishing the two genotype categories. Relative change in traits in response to soil environments was computed for each genotype and timepoint using the following formula: (*trait*_*CL*_ — *trait*_*SL*_)/*trait*_*avg*_, where *trait*_*CL*_ indicates the trait value under clay loam conditions, and *trait*_*SL*_ indicates the trait value under sandy loam conditions, and *trait*_*avg*_ indicates the average trait value under both conditions. Heatmaps of Pearson correlations of relative changes between shoot bioimass and all root traits were generated using the *corrplot* R package (Wei et al., 2017).

To estimate the effect of soil environment (sandy loam or clay loam), genotype group (breeding line or diversity line), and their interactions on root traits observed by 2D imaging, a linear mixed effects model was fit using the ‘lmer’ function from the *lme4* package in R (Bates, 2010). The three main assumptions (linearity, normality of residuals and homoscedasticity) were tested and found to be met. The model was specified as follows:

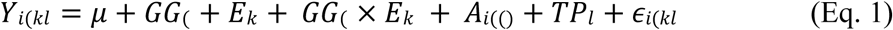

In Equation 1, *Y*_*i*(*kl*_ is the trait value, μ is the mean, *GG*(is the fixed effect of genotype group *j*, 𝐸𝑘is the fixed effect of the environment *k*, 𝐺𝐺(× 𝐸𝑘 is the interaction effect of genotype group *j* and environment *k*, 𝐴𝑖(() is the random effect of genotype *i* nested within genotype group *j*, 𝑇𝑃𝑙 is the random effect of timepoint *l*, and 𝜖𝑖(𝑘𝑙 is the residual error. In this model, the fixed effects were environment, genotype group, and their interactions. To quantify potential interactions between environment and specific genotypes across different sampling timepoints, we fit additional linear models for each of the three timepoints. One complicating factor for studies on soybean root variation is that root development is linked to plant growth stages, with distinct patterns emerging throughout the plant’s lifecycle. Thus, the timepoints were considered separately. The model was specified as follows:

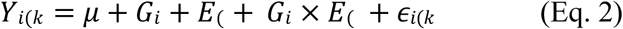

In Equation 2, 𝑌𝑖(𝑘is the trait value, 𝜇 is the mean, 𝐺𝑖is the fixed effect of genotype *i*, 𝐸(is the fixed effect of the environment *j*, 𝐺𝑖 × 𝐸(is the interaction effect of genotype *i* and environment *j*, and 𝜖𝑖(𝑘 is the residual error. Broad sense heritability for each root trait was estimated using linear mixed models fitted with the R-package *lme4* (Bates, 2010). Genotype was treated as a random effect and variance components were used to calculate broad sense heritability. Narrow sense heritability was estimated using the *sommer* package (Covarrubias-Pazaran, 2025). Models were fitted using the “mmes” function with genotype modeled as a random effect and the genomic kinship matrix specified via the “vsm” function. Narrow sense heritability was calculated as the ratio of additive genetic variance to the total phenotypic variance, following:

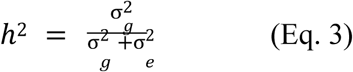

where ℎ^2^ is narrow sense heritability, 𝜎^2^ is the additive genetic variance, and 𝜎^2^ is the residual variance.

### Data availability

All data associated with this study are available on Zenodo at doi: 10.5281/zenodo.11037393

## Results and Discussion

### Genetic differentiation of genotype groups

Results from PCA and hierarchical clustering using 6K genome-wide SNPs supported the strong differentiation of lines according to their originating source: the breeder lines developed for Midwest United States high input production environments (BL group) versus diversity lines selected from the USDA Soybean Germplasm Collection (DL group), which are considered landraces (**Figure S4**). Interestingly, there was one U.S. line (cv. Lincoln) from the DL group which, despite being developed for production in the Midwest U.S. by Illinois Agricultural Experimental Station (Weiss, 1953), grouped along with the rest of the diversity lines. Lincoln was developed over 70 years ago and subsequently used as the parent in other crosses. For our study, this line serves as a baseline of comparison between historical U.S. varieties and the modern elite lines of the BL group. PC1 of the PCA explained 38.9% of total variance and clearly separated the BL and DL. PC2 explained 10.7% of total variance and accounted for within-DL variation. The cultivar, Lincoln, was the closest diversity line to the BL group along the PC1 axis and mapped in line with the BL along the PC2 axis, consistent with its U.S. origins. Having observed this strong differentiation between the genotype groups of breeding lines and diversity lines, we next examined root trait variation as it related to genotype group.

### Trait relationships are affected by growth stage and genotype group

Correlation matrices of the 12 RSA traits, root biomass, and shoot biomass were analyzed for the entire dataset as well as for data subsets divided by timepoints and genotype groups. Positive correlations were prevalent in the overall dataset (**Figure S5a**). However, this was likely due to high levels of root system growth across developmental time, especially between TP1 and TP2, leading to larger values of all traits across timepoints, resulting in positive correlations throughout and potentially masking some true negative correlations. Indeed, when data were separated by timepoints, both positive and negative correlations emerged (**Figure S5b-d**). For example, negative correlations were found between root diameter and number of forks (r = -0.32) and number of tips (r = -0.52), suggesting a trade-off between root diameter and branching structures consistent with previous studies (Eissenstat et al., 2015; Comas and Eissenstat, 2009). Positive correlations, such as those between root biomass and RSA traits (root volume, root surface area, and root diameter), were observed to increase throughout development. Nodule count was generally uncorrelated with RSA or biomass traits, except for a weak negative correlation with average root angle during the earliest timepoint (r = -0.25). This result sits in contrast with earlier reports of positive correlations between nodules and shoot/root weight at the R3 developmental stage (Sinclair et al., 1991).

Quantifying the similarity between trait relationships across time, Spearman’s correlation, rs, between matrix pairs showed that relationships in TP2 and TP3 were more similar to each other (r = 0.88) than either of those timepoints were with TP1 (r = 0.76 between TP1 and TP2; r = 0.78 between TP1 and TP3). This could be due to the transition between vegetative and reproductive phase that occurred around TP2, whereas TP1 represented early vegetative growth. Comparing correlation matrices between breeder lines and diversity lines across timepoints, trait correlations tended to be stronger (in both negative and positive directions) in the breeder line group (**Figure S6**). Spearman’s correlation of the matrices across timepoints showed higher values (rs values of 0.57, 059, and 0.65 for timepoint pairs) compared to the diversity line group (rs values of 0.48, 0.31, and 0.58 for the same timepoint pairs).

### Root trait variation can differentiate genotype groups

Principal component analysis of trait data, averaged across replicates and environments, revealed that PCs 1 and 2 together accounted for ∼80% of total variation (PC1: 64.22 %; PC2: 15.05%) (**Figure 2a**).

**Figure 2.**
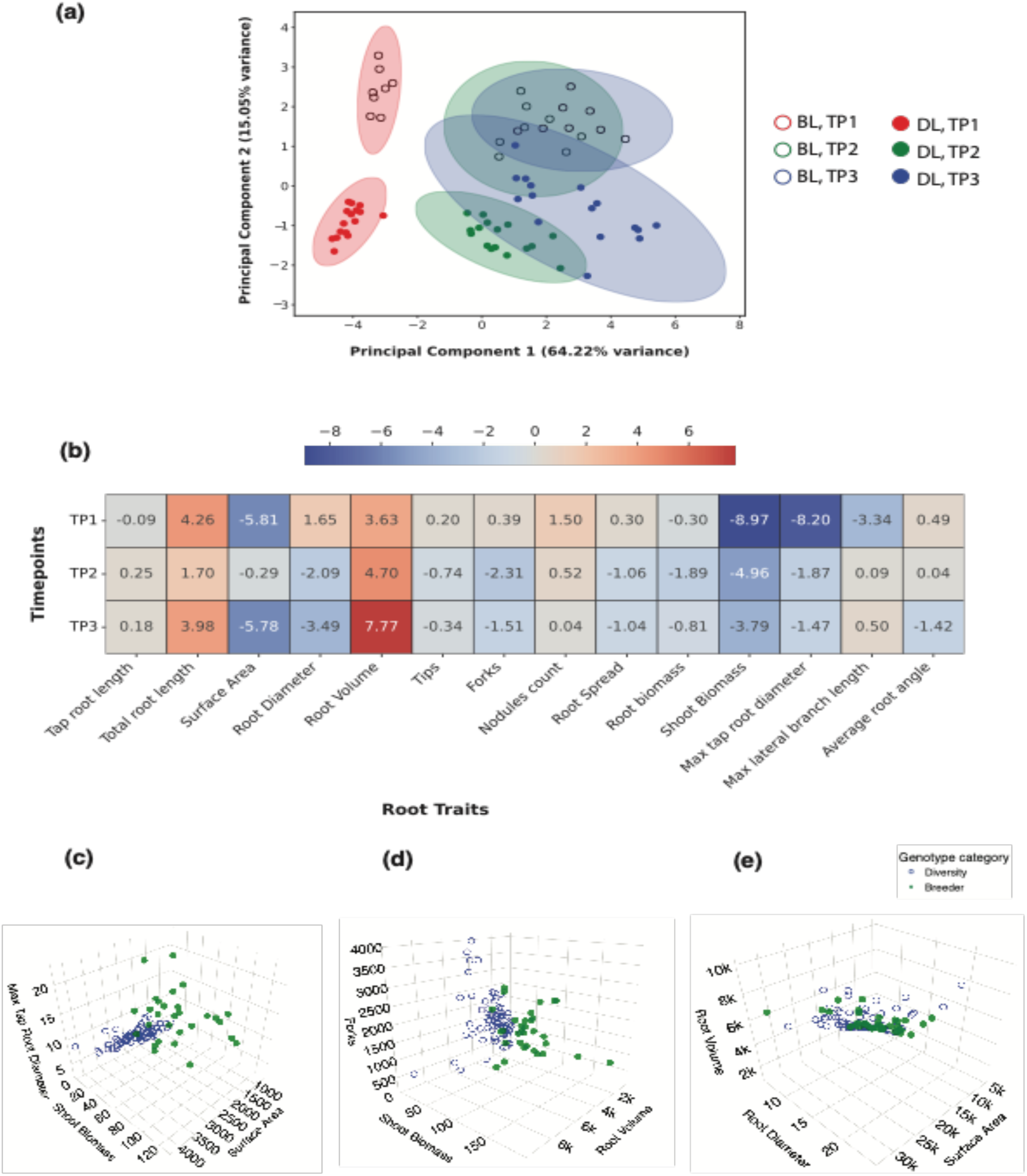
Differentiation of genotype groups by root traits and shoot biomass: **(a)** Results of Principal Components Analysis of genotype means (averaged across environments and replicates) colored by their sampling timepoints (TP1, TP2, and TP3) and genotype group (BL: breeding line; DL: diversity line). (b) Heatmap showing coefficients of 13 root traits along with shoot biomass resulting from Linear Discriminant Analysis to distinguish genotype groups. 3D Scatter plot comparing diversity lines and breeder lines of (c) maximum tap root diameter, surface area and shoot biomass, (d) shoot biomass, number of forks and root volume, and (e) root volume, surface area and average root diameter.

Evaluating the loadings of each trait variable, we observed that traits such as surface area (loading = 0.98), root volume (0.96), and total root length (0.95) were the top contributors to PC1. With regard to PC2, shoot biomass (0.83), maximum tap root diameter (0.77), and maximum lateral branch length (0.66) were the main contributors. PCA additionally revealed clusters that corresponded to specific combinations of genotype group and timepoint, suggesting there were stark phenotypic differences among these groups; PC1 was aligned with the sample timepoint while PC2 could be explained by genotype group (BL versus DL). When comparing observations from TP1 versus TP2 and TP3, we observed that TP1 formed a tighter cluster and was strongly separated from the other two groups, consistent with the greater similarity between TP2 and TP3 trait relationships described earlier. The TP1 sampling corresponded to early vegetative development (V2-V6), and as plants matured in TP2 and TP3, their traits may have had increased influence from interactions between the genotype and the temporally varying environment, leading to differential resource allocations and more variation among observations (Sultan, 2000; Gratani, 2014). Comparing clusters of BLs versus those formed by DLs, we noted that observations from the BL group were less dispersed than those from the DL group in the PC1-PC2 space. This could be a result of decreased genetic variation in breeding germplasm as compared to landraces (Hyten et al., 2006). Alternatively, this may reflect the greater number of diversity lines evaluated in our panel.

We next asked which traits jointly contributed the most to the differentiation between DL and BL genotype groups. Fisher’s LDA coefficients indicated that the top contributing traits were generally consistent across timepoint (**Figure 2b-e**). For example, shoot biomass, total root length, and root volume ranked in the top five predictors across all three timepoints. In addition to those three traits, other strong contributors to the discriminant score were maximum taproot diameter and surface area (in TP1), number of forks and root biomass (in TP2), and surface area and average root diameter (in TP3). Although previous work on root trait variation in a large collection of soybean germplasm, including elite cultivars and landraces, also found differentiation among these groups, the traits likely responsible were not examined (Prince et al., 2020).

### Evidence for greater root plasticity in response to different soil environments in diverse soybean lines

To quantify the effects of genotype group, soil environment and their interaction on individual traits, we fitted a mixed linear model with timepoint as a random variable (**Methods**). The main effect of genotype group was not significant for any trait examined but the main effect of environment was significant (p<0.01) for five traits (tap root length, root volume, root spread, root biomass, and average root diameter). In clay loam soil, each of these traits showed reductions compared to sandy loam soil (tap root length by 14.20%; root biomass by 18%; root volume by 22.88%; root spread by 22.73%; and average root diameter by 7.55%). We speculate two different reasons for these results. One, the heavier textured clay loam soil could have been more difficult for roots to penetrate as compared to the looser structure found in the sandy loam environment (Cairns et al., 2011). Two, explanation for these root trait differences could be that the lower nutrient and water holding capacity of the sandier soils may have necessitated greater allocation of resources to root growth for increased foraging than in the clay loam; results from soil analysis showed the sandy loam environment had lower levels of calcium, cation exchange capacity, and water holding capacity but higher levels of manganese (**Figure S1**).

Three traits (maximum lateral branch length, maximum tap root diameter, and number of tips) showed a significant effects of the interaction between genotype group and environment, indicating that the two genotype groups showed a differential response to changes in soil type. Interestingly, in all three cases, the group composed of diversity lines was found to show a much stronger response to the environment (**Figure 3**).

**Figure 3.**
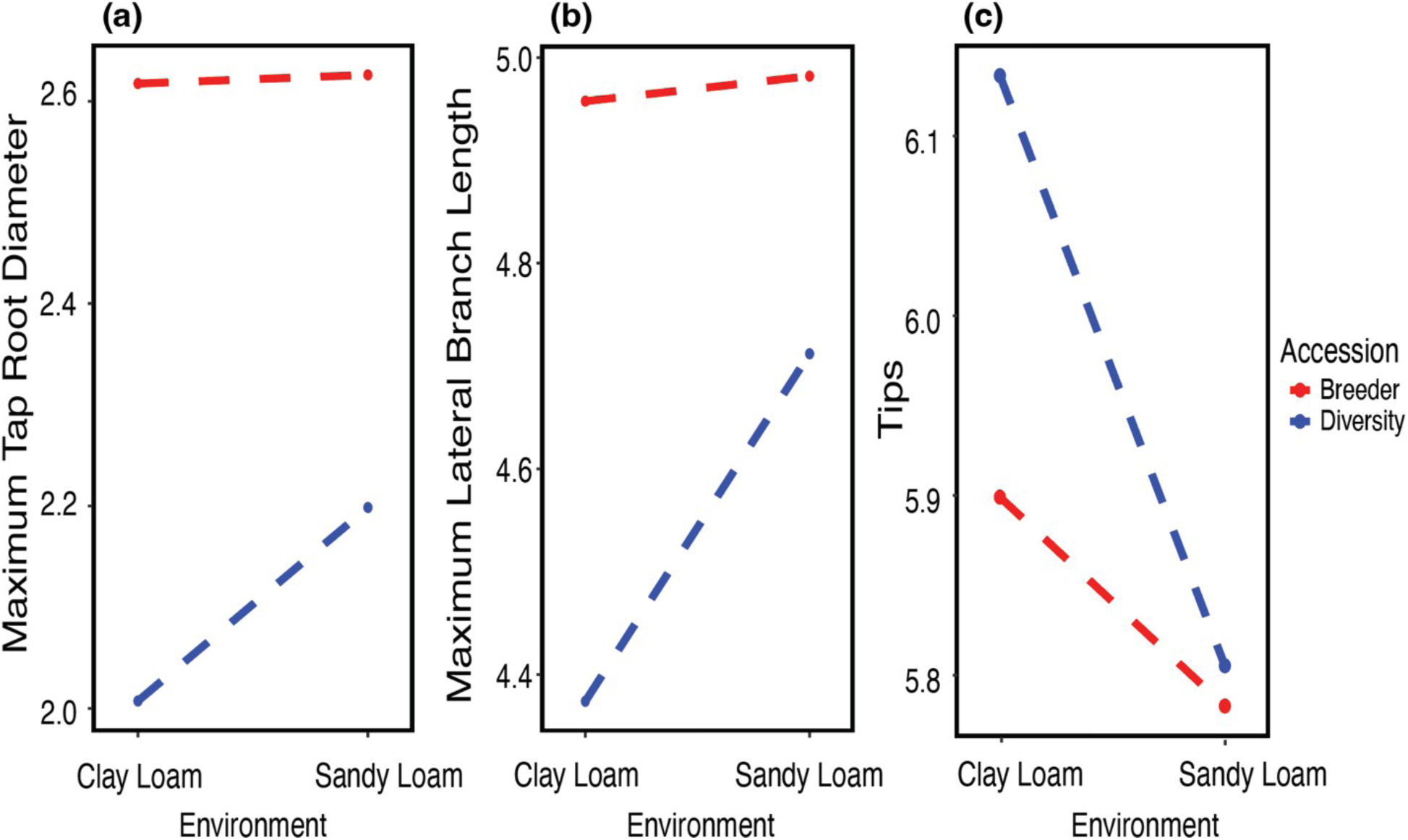
Interaction plot of root traits across soil types for genotype groups. The transformed mean trait values are shown for two soil environments (Clay Loam and Sandy Loam) in breeder (red dashed line) and diversity (blue dashed line) lines. **(a)** Maximum lateral branch length, **(b)** Maximum tap root diameter, **(c)** Number of root tips. Mixed model results are shown in Table 2.

**Table 2.** Summary of linear mixed effects model results. Significance of genotype group, environment, and their interaction resulting from linear mixed effects models (Equation 1) fit for each of the 14 root traits along with shoot biomass. Significance levels: ’***’ : p <0.001, ’**’: p <0.01, ’*’: p < 0.05, ’.’: p < 0.1, ns : not significant

| Trait | GG | E | GG × E |
| --- | --- | --- | --- |
| Average root angle | ns | ns | ns |
| Average root diameter | ns | ** | ns |
| Forks | ns | ns | ns |
| Maximum lateral branch length | ns | ns | ** |
| Maximum tap root diameter | ns | ns | ** |
| Nodules | ns | ns | ns |
| Root biomass | ns | ** | ns |
| Root spread | ns | *** | ns |
| Root volume | ns | ** | ns |
| Shoot biomass | ns | ns | ns |
| Surface area | ns | ns | ns |
| Tap root length | ns | ** | ns |
| Tips | ns | ns | * |
| Total root length | ns | ns | ns |

For maximum lateral branch length, DL was 7.72% longer in sandy loam than in clay loam soil, whereas BL was barely affected at 0.49%. Similarly, for maximum tap root diameter, the DL was 9.52% greater in sandy loam than in clay loam soil, whereas BL was only affected by 0.32%. Finally, for the number of tips, the DL had 5.25% reduction in the number of tips in the sandy loam soil than in the clay loam soil; meanwhile, the BL showed a 1.86% reduction under sandy loam soil. Taken together, these results indicate greater phenotypic plasticity in root architecture among diverse lines of soybean compared to elite breeding material. In contrast to the significant increases for size traits (*i.e.* length, volume, mass) under sandy loam soil, the number of tips, which is a count trait, was observed to decrease in both genotype groups, with a stronger response for DL than BL (p<0.05). As mentioned previously, in heavier textured soils with greater proportions of fine particles, there is greater resistance to root penetration, possibly causing more lateral branching. This notion is supported by previous findings that sandier soils with larger particles enable greater aeration and lower mechanical impedance (Comas et al., 2013).

Beyond high-level genotype group classifications, we tested for genotype by environment interactions on root traits and shoot biomass (Table 2). Separate models were fit for each timepoint and included genotype and environment main effects and their interaction. (**Figure 4** and **Table S3**). In contrast, many traits were significant for Genotype main effect only, and even greater numbers of traits showed significant G × E effects. Interestingly, there was a distinct inverse relationship between the numbers of traits that had significant G × E and the numbers of traits that had significant Genotype main effect only; G × E became more prevalent towards later stages of development while G main effects decreased (**Figure 4**). Traits where G x E effects were significant across all three timepoints were maximum tap root diameter, nodule number, root biomass, and root spread.

**Figure 4.**
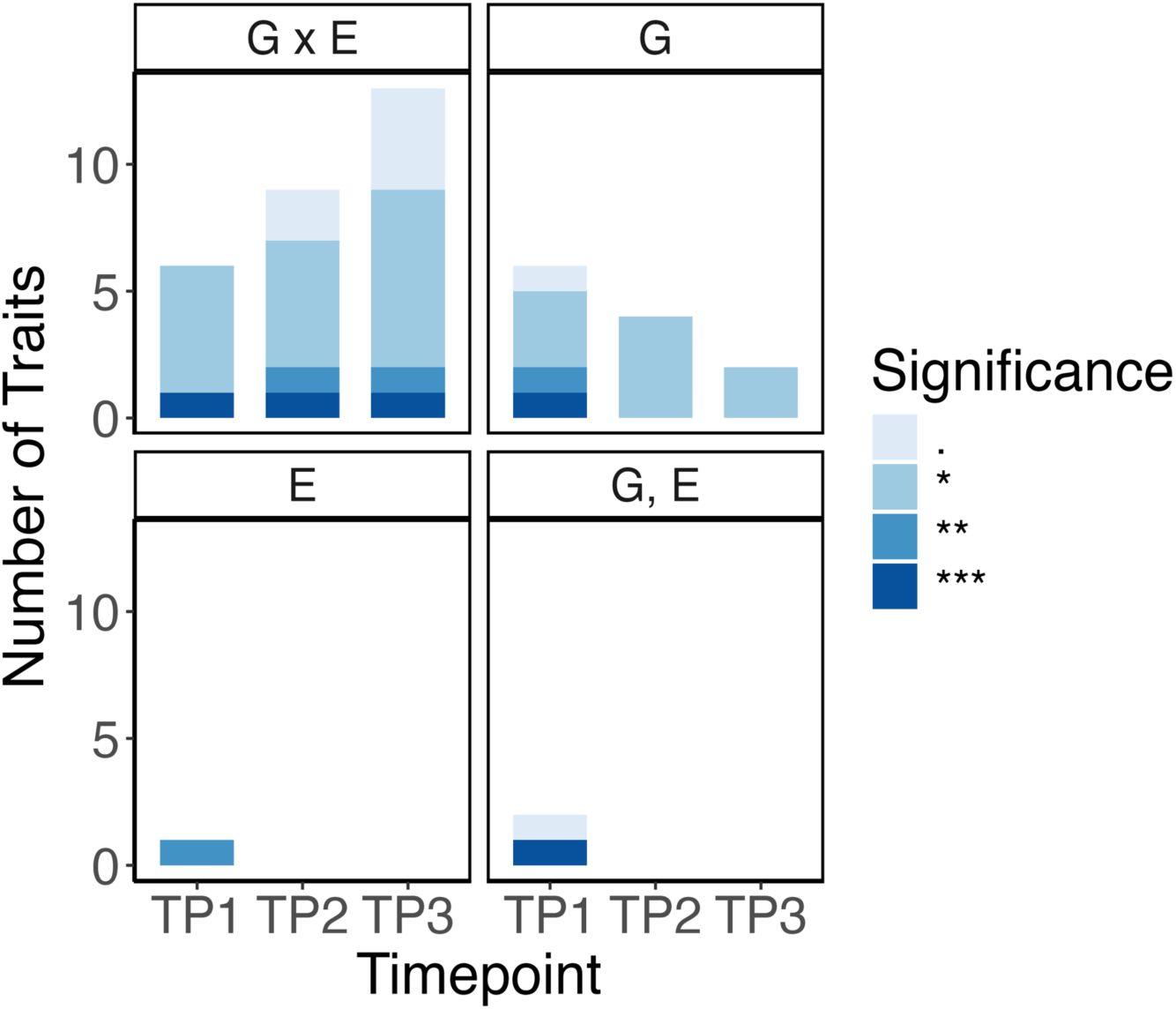
Effects of genotype, environment, and genotype x environment interaction on root traits across timepoints. Results of models shown in **Table S3** are summarized here for the total number of traits that had **(a)** significant G x E interaction effects; **(b)** only significant Genotype main effects; **(c)** only significant Environment main effects; and **(d)** significant Genotype and Environment main effects but no significant G x E. Models were fit separately for each timepoint. Significance levels: ’***’: p <0.001, ’**’: p <0.01, ’*’: p < 0.05, ’.’: p < 0.1

Finally, we next asked whether belowground phenotypic plasticity corresponded to plasticity aboveground. Phenotypic plasticity was represented by the relative change in trait values between the two soil environments (see **Methods** for calculation). We found that negative correlations were prevalent between belowground traits and shoot biomass with respect to plasticity, except for in timepoint two where positive correlations emerged (**Figure S7**). Most notably, relative changes to maximum taproot diameter and maximum lateral branch length were positively correlated with relative change to shoot biomass during this transition to reproductive growth. Overall results from timepoints one and three suggest a tradeoff in aboveground and belowground trait plasticity in response to soil environment. Divergence of timepoint two results from the earlier and later developmental stages could be due to this sampling occurring around the transition period from vegetative to reproductive growth, which involves major changes to plant physiology.

### Maximum taproot diameter and maximum lateral branch length have the highest heritability among measured root traits

We next evaluated broad- and narrow-sense heritabilities of root traits across the two environments and three growth stages. Across timepoints, heritabilities were highest in TP1. However, consistent with previous studies, our root measures showed generally low heritabilities (**Figure S8**). Since differences in mean heritabilities between environments were not significant, we averaged heritabilities across environments for further analysis. Heritabilities across timepoints were highest in TP1. Overall mean values were highest for maximum taproot diameter (*H^2^* = 0.37 and *h^2^* = 0.21) and maximum lateral branch length (*H^2^* = 0.22 and *h^2^* = 0.13) among root traits. These values were lower than heritabilities computed for shoot biomass (*H^2^* = 0.57 and *h^2^* = 0.24), which served as a benchmark as the sole aboveground trait. Heritabilities for maximum taproot diameter and maximum lateral branch length were actually comparable (or slightly higher) than those for shoot biomass during TP1 but decreased linearly over the subsequent timepoints. In contrast, shoot biomass remained more consistent (**Figure 5**).

**Figure 5.**
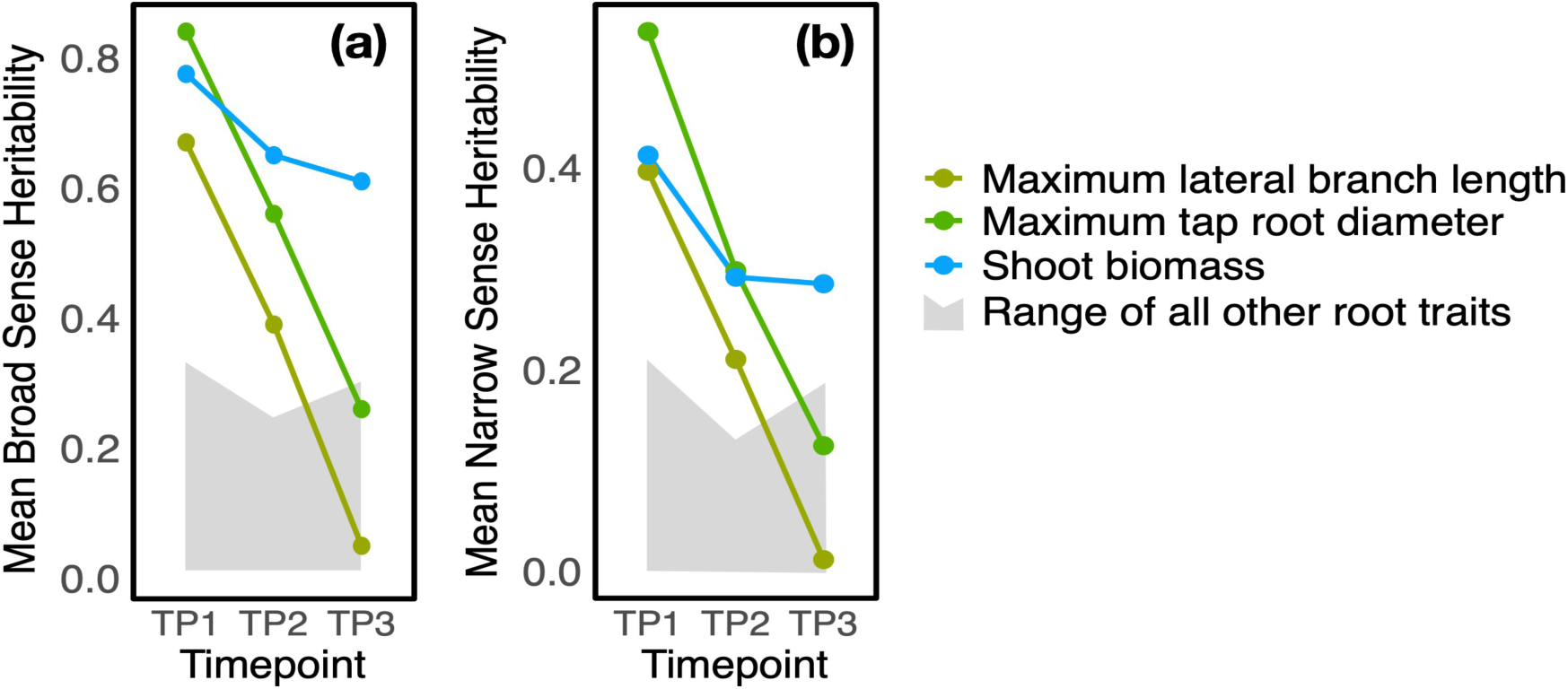
Trait heritability across timepoints. **(a)** Mean broad-sense heritability (H²) and **(b)** mean narrow-sense heritability (h²) for selected traits across three timepoints (TP1–TP3). Colored lines represent three representative traits: maximum lateral branch length (olive), maximum tap root diameter (green), and shoot biomass (blue). The shaded gray area indicates the range of heritability values for all other root traits.

The observed reductions in heritability over time were unique to these two RSA traits: other root traits showed overall lower but more consistent heritability values across timepoints (shaded polygon in **Figure 5**). Interestingly, both these traits were extracted by first making four to five random measurements manually per 2D image, representing only a small sample of what is found within a single root system. The dramatic decrease in heritabilities across development could potentially be explained by increased error in our 2D measurement approach as root systems grew larger from TP1 to TP2 and TP3.

### Characterizing root trait distributions using a portable, low-cost 3D method

Methods that can better describe trait distributions in root systems may reduce measurement error, particularly for traits requiring multiple measurements within a single root system, such as maximum tap root diameter and maximum lateral branch length. We tested the application of 3D root modeling to extract features using a new open-source framework. A total of 39 traits were extracted on a subset of 30 root systems as a proof-of-concept (**Methods)**. Of these traits, 12 were also measured using the 2D approach. In comparing trait values estimated from 2D versus 3D methodologies, 5 out of the 12 root traits showed correlations (*r* = 0.6–0.8) in measurements between the two methods (**Figure S9**). For traits such as lateral branch length or branching angle, which has a distribution for each root system, 3D models enabled detection of values across a much larger range. For example, comparisons of 2D and 3D distributions for lateral branch length demonstrated that 3D feature extraction enabled sampling of the shortest branches, whereas the 2D method relied on human-selection of “random” branches, potentially biasing selections towards longer branches (**Figures S10-S15**). This seems especially likely since much of the distribution of lateral branch lengths lies in the lower-value range. Moreover, a study comparing 2D and 3D methodologies found that 3D methods were better for estimating lateral branch lengths in the tree species, *Pinus pinaste*r (Danjon et al., 1999). Nevertheless, simpler traits (e.g., root length and diameter) are still adequately extracted using 2D methods (Clark et al., 2013). Perhaps the greatest advantage of root trait estimation using 3D models is that this approach can enable extraction of novel RSA descriptors (e.g., minimum, maximum, or coefficient of variation for root trait distributions) for use in downstream analyses such as genetic mapping.

## Conclusion

While breeding programs have been historically successful in developing new soybean cultivars, there is an increasingly narrow genetic base within breeder panels (Moose & Mumm, 2008). Diversity panels of landraces or wild accessions represent a source of novel alleles offering higher levels of genetic and adaptive potential to changing environments (Meyer et al. 2012; Wei & Jiang, 2021), particularly in the case of future challenging climatic scenarios. Our study finds evidence for divergence of root system traits in cultivated soybean as an indirect result of modern breeding. In the context of our study, the root systems of elite breeding lines were larger but less plastic in response to changing soil environments than those of landraces. These findings contrast with previous conclusions that more parsimonious root systems underlie superior biomass in elite lines (Noh et al., 2022). Rarther they are more aligned with results from an era study suggesting that soybean root systems became larger over time as a result of selection (Mandozai et al., 2021). While heritabilities of root traits are generally low, the higher values for traits like maximum tap root diameter and maximum lateral branch length suggest that using measurement approaches that more fully characterize within-sample trait distributions can help decrease measurement error and improve heritability estimates. Future advancements in 3D root modeling and feature extraction could help to better quantify RSA towards more translational opportunities for root architecture traits in varietal crop improvement.

## Acknowledgements

The authors gratefully acknowledge Karla Miserendino, Andrea Portillo, Catherine Mejia, To- Chia Ting, Luis Vargas, Sajad Jamshidi, Ava Antic, Henrique Feiler, Chancelor Clark, and Weidong Wang for technical assistance; Jim Camberato for guidance on soil analysis; and Katy Rainey for providing seed from the Public Biomass Panel. This work was funded by grants USDA AFRI #2022-67013-36205 and NSF-PGRP #2102120 to DRW and by Purdue University as part of AgSEED Crossroads funding to support Indiana’s Agriculture and Rural Development.

## References

1. Atkinson, J. A., Pound, M. P., Bennett, M. J., & Wells, D. M. (2019). Uncovering the hidden half of plants using new advances in root phenotyping. Current opinion in biotechnology, 55, 1–8.

2. Bates, D. M. (2010). lme4: Mixed-effects modeling with R.

3. Bradshaw, A. D. (2006). Unravelling phenotypic plasticity–why should we bother? New Phytologist, 170(4), 644–648.

4. Cairns, J. E., Impa, S. M., O’Toole, J. C., Jagadish, S. V. K., & Price, A. H. (2011). Influence of the soil physical environment on rice (Oryza sativa L.) response to drought stress and its implications for drought research. Field Crops Research, 121(3), 303–310.

5. Clark, R. T., Famoso, A. N., Zhao, K., Shaff, J. E., Craft, E. J., Bustamante, C. D., Mccouch,S.R, Aneshansley. D.J., & Kochian, L. V. (2013). High-throughput two-dimensional root system phenotyping platform facilitates genetic analysis of root growth and development. Plant, cell & environment, 36(2), 454–466.

6. Comas, L. H., & Eissenstat, D. M. (2009). Patterns in root trait variation among 25 co-existing North American forest species. New Phytologist, 182(4), 919–928.

7. Comas, L. H., Becker, S. R., Cruz, V. M. V., Byrne, P. F., & Dierig, D. A. (2013). Root traits contributing to plant productivity under drought. Frontiers in plant science, 4, 442.

8. Covarrubias-Pazaran, G., & Covarrubias-Pazaran, M. G. (2025). Package ‘sommer’.

9. Danjon, F., Bert, D., Godin, C., & Trichet, P. (1999). Structural root architecture of 5-year-old Pinus pinaster measured by 3D digitising and analysed with AMAPmod. Plant and Soil, 217(1), 49–63.

10. Eissenstat, D. M., Kucharski, J. M., Zadworny, M., Adams, T. S., & Koide, R. T. (2015). Linking root traits to nutrient foraging in arbuscular mycorrhizal trees in a temperate forest. New Phytologist, 208(1), 114–124.

11. Fradgley, N., Evans, G., Biernaskie, J.M., Cockram, J., Marr, E.C., Oliver, A.G., Ober, E. & Jones, H., (2020). Effects of breeding history and crop management on the root architecture of wheat. Plant and Soil, 452, pp.587–600.

12. Ghalambor, C. K., McKay, J. K., Carroll, S. P., & Reznick, D. N. (2007). Adaptive versus non-adaptive phenotypic plasticity and the potential for contemporary adaptation in new environments. Functional ecology, 21(3), 394–407.

13. Gratani, L. (2014). Plant phenotypic plasticity in response to environmental factors. Advances in botany, 2014(1), 208747.

14. Hoogenboom, G., Huck, M. G., & Peterson, C. M. (1987). Root growth rate of soybean as affected by drought stress 1. Agronomy Journal, 79(4), 607–614.

15. Hunter, J., & Dale, D. (2007). The matplotlib user’s guide. Matplotlib 0.90. 0 user’s guide, 487.

16. Hyten, D.L., Song, Q., Zhu, Y., Choi, I.Y., Nelson, R.L., Costa, J.M., Specht, J.E., Shoemaker, R.C. and Cregan, P.B., 2006. Impacts of genetic bottlenecks on soybean genome diversity. Proceedings of the National Academy of Sciences, 103(45), pp.16666–16671.

17. Kuijken, R. C., van Eeuwijk, F. A., Marcelis, L. F., & Bouwmeester, H. J. (2015). Root phenotyping: from component trait in the lab to breeding. Journal of experimental botany, 66(18), 5389–5401.

18. Lê, S., Josse, J., & Husson, F. (2008). FactoMineR: an R package for multivariate analysis. Journal of statistical software, 25, 1–18.

19. Lynch, J. P. (2019). Root phenotypes for improved nutrient capture: an underexploited opportunity for global agriculture. New phytologist, 223(2), 548–564.

20. Lynch, J. P., Mooney, S. J., Strock, C. F., & Schneider, H. M. (2022). Future roots for future soils. Plant, Cell & Environment, 45(3), 620–636.

21. Malamy, J. E. (2005). Intrinsic and environmental response pathways that regulate root system architecture. Plant, cell & environment, 28(1), 67–77.

22. Mandozai, A., Moussa, A.A., Zhang, Q., Qu, J., Du, Y., Anwari, G., Al Amin, N. and Wang, P., 2021. Genome-wide association study of root and shoot related traits in spring soybean (Glycine max L.) at seedling stages using SLAF-seq. Frontiers in Plant Science, 12, p.568995.

23. McNear Jr, D. H. (2013). The rhizosphere-roots, soil and everything in between. Nature Education Knowledge, 4(3), 1.

24. Messina, C., McDonald, D., Poffenbarger, H., Clark, R., Salinas, A., Fang, Y., Gho, C., Tang, T., Graham, G., Hammer, G.L. and Cooper, M., 2021. Reproductive resilience but not root architecture underpins yield improvement under drought in maize. Journal of Experimental Botany, 72(14), pp.5235–5245.

25. Meyer, R. S., DuVal, A. E., & Jensen, H. R. (2012). Patterns and processes in crop domestication: an historical review and quantitative analysis of 203 global food crops. New Phytologist, 196(1), 29–48.

26. Moose, S. P., & Mumm, R. H. (2008). Molecular plant breeding as the foundation for 21st century crop improvement. Plant physiology, 147(3), 969–977.

27. Noh, E., Fallen, B., Payero, J., & Narayanan, S. (2022). Parsimonious root systems and better root distribution can improve biomass production and yield of soybean. Plos one, 17(6), e0270109.

28. O’toole, J. C., & Bland, W. L. (1987). Genotypic variation in crop plant root systems. Advances in agronomy, 41, 91–145.

29. Ordonez, R. A., Archontoulis, S. V., Martinez-Feria, R., Hatfield, J. L., Wright, E. E., & Castellano, M. J. (2020). Root to shoot and carbon to nitrogen ratios of maize and soybean crops in the US Midwest. European Journal of Agronomy, 120, 126130.

30. Pedregosa, F., Varoquaux, G., Gramfort, A., Michel, V., Thirion, B., Grisel, O., Blondel, M., Prettenhofer, P., Weiss, R., Dubourg, V. & Vanderplas, J. (2011). Scikit-learn: Machine learning in Python. the Journal of machine Learning research, 12, pp.2825–2830.

31. Prince, S.J., Vuong, T.D., Wu, X., Bai, Y., Lu, F., Kumpatla, S.P., Valliyodan, B., Shannon, J.G. and Nguyen, H.T. (2020). Mapping quantitative trait loci for soybean seedling shoot and root architecture traits in an inter-specific genetic population. Frontiers in Plant Science, 11, p.1284.

32. Rinehart, B., Borras, L., Salmeron, M., McNear Jr, D. H., & Poffenbarger, H. (2024). Commercial maize hybrids have smaller root systems after 80 years of breeding. Rhizosphere, 30, 100915.

33. Robinson, D. (2001). Root proliferation, nitrate inflow and their carbon costs during nitrogen capture by competing plants in patchy soil. Plant and Soil, 232, 41–50.

34. Schneider, C. A., Rasband, W. S., & Eliceiri, K. W. (2012). NIH Image to ImageJ: 25 years of image analysis. Nature methods, 9(7), 671–675.

35. Schneider, H. M., & Lynch, J. P. (2020). Should root plasticity be a crop breeding target? Frontiers in Plant Science, 11, 546.

36. Sinclair, T. R., Soffes, A. R., Hinson, K., Albrecht, S. L., & Pfahler, P. L. (1991). Genotypic variation in soybean nodule number and weight. Crop science, 31(2), 301–304.

37. Song, P., Wang, J., Guo, X., Yang, W., & Zhao, C. (2021). High-throughput phenotyping: Breaking through the bottleneck in future crop breeding. The Crop Journal, 9(3), 633–645.

38. Song, Q., Hyten, D.L., Jia, G., Quigley, C.V., Fickus, E.W., Nelson, R.L. and Cregan, P.B., 2013. Development and evaluation of SoySNP50K, a high-density genotyping array for soybean. PloS one, 8(1), p.e54985._

39. Song, Q., Hyten, D.L., Jia, G., Quigley, C.V., Fickus, E.W., Nelson, R.L. and Cregan, P.B., 2015. Fingerprinting soybean germplasm and its utility in genomic research. G3: Genes, genomes, genetics, 5(10), pp.1999-2006.

40. Stingaciu, L., Schulz, H., Pohlmeier, A., Behnke, S., Zilken, H., Javaux, M., & Vereecken, H. (2013). In situ root system architecture extraction from magnetic resonance imaging for water uptake modeling. Vadose zone journal, 12(1), vzj2012-0019.

41. Sultan, S. E. (2000). Phenotypic plasticity for plant development, function and life history. Trends in Plant Science, 5(12), 537–542.

42. Sultan, S. E. (2015). Organism and environment: ecological development, niche construction, and adaptation. Oxford University Press.

43. Tracy, S. R., Nagel, K. A., Postma, J. A., Fassbender, H., Wasson, A., & Watt, M. (2020). Crop improvement from phenotyping roots: highlights reveal expanding opportunities. Trends in plant science, 25(1), 105–118.

44. Tuberosa R, Salvi S, Sanguineti MC, Landi P, Maccaferri M, Conti S (2002) Mapping QTLs regulating morpho-physiological traits and yield: case studies, shortcomings and perspectives in drought stressed maize. Annals of Botany, 89:941–963 10.1093/aob/mcf134

45. Uga, Y., Sugimoto, K., Ogawa, S., Rane, J., Ishitani, M., Hara, N., Kitomi, Y., Inukai, Y., Ono, K., Kanno, N. and Inoue, H. (2013). Control of root system architecture by DEEPER ROOTING 1 increases rice yield under drought conditions. Nature genetics, 45(9), pp.1097–1102.

46. Van Kleunen, M., & Fischer, M. (2005). Constraints on the evolution of adaptive phenotypic plasticity in plants. New phytologist, 166(1), 49–60.

47. Van Rossum, G., & Drake, F. L. (2009). Python/C Api manual-python 3. CreateSpace.

48. Wasson, A.P., Richards, R.A., Chatrath, R., Misra, S.C., Prasad, S.S., Rebetzke, G.J., Kirkegaard, J.A., Christopher, J. and Watt, M. (2012). Traits and selection strategies to improve root systems and water uptake in water-limited wheat crops. Journal of experimental botany, 63(9), pp.3485–3498.

49. Watt, M., Silk, W. K., & Passioura, J. B. (2006). Rates of root and organism growth, soil conditions, and temporal and spatial development of the rhizosphere. Annals of botany, 97(5), 839–855.

50. Wei, T., Simko, V., Levy, M., Xie, Y., Jin, Y., & Zemla, J. (2017). Package ‘corrplot’. Statistician, 56(316), e24

51. Wei, X., & Jiang, M. (2021). Meta-analysis of genetic representativeness of plant populations under ex situ conservation in contrast to wild source populations. Conservation Biology, 35(1), 12–23.

52. Weiss, M.G. (1953). Registration of Soybean Varieties, III. Agronomy Journal 45, 326–330.

53. Wickham, H. (2016). Getting Started with ggplot2. In ggplot2: Elegant graphics for data analysis (pp. 11–31). Cham: Springer International Publishing.

54. York, L. M., & Lynch, J. P. (2015). Intensive field phenotyping of maize (Zea mays L.) root crowns identifies phenes and phene integration associated with plant growth and nitrogen acquisition. Journal of Experimental Botany, 66(18), 5493–5505.

