## Supplementary material for "Divergence of root system traits in soybean between breeding and diversity lines": Supplimental Figures

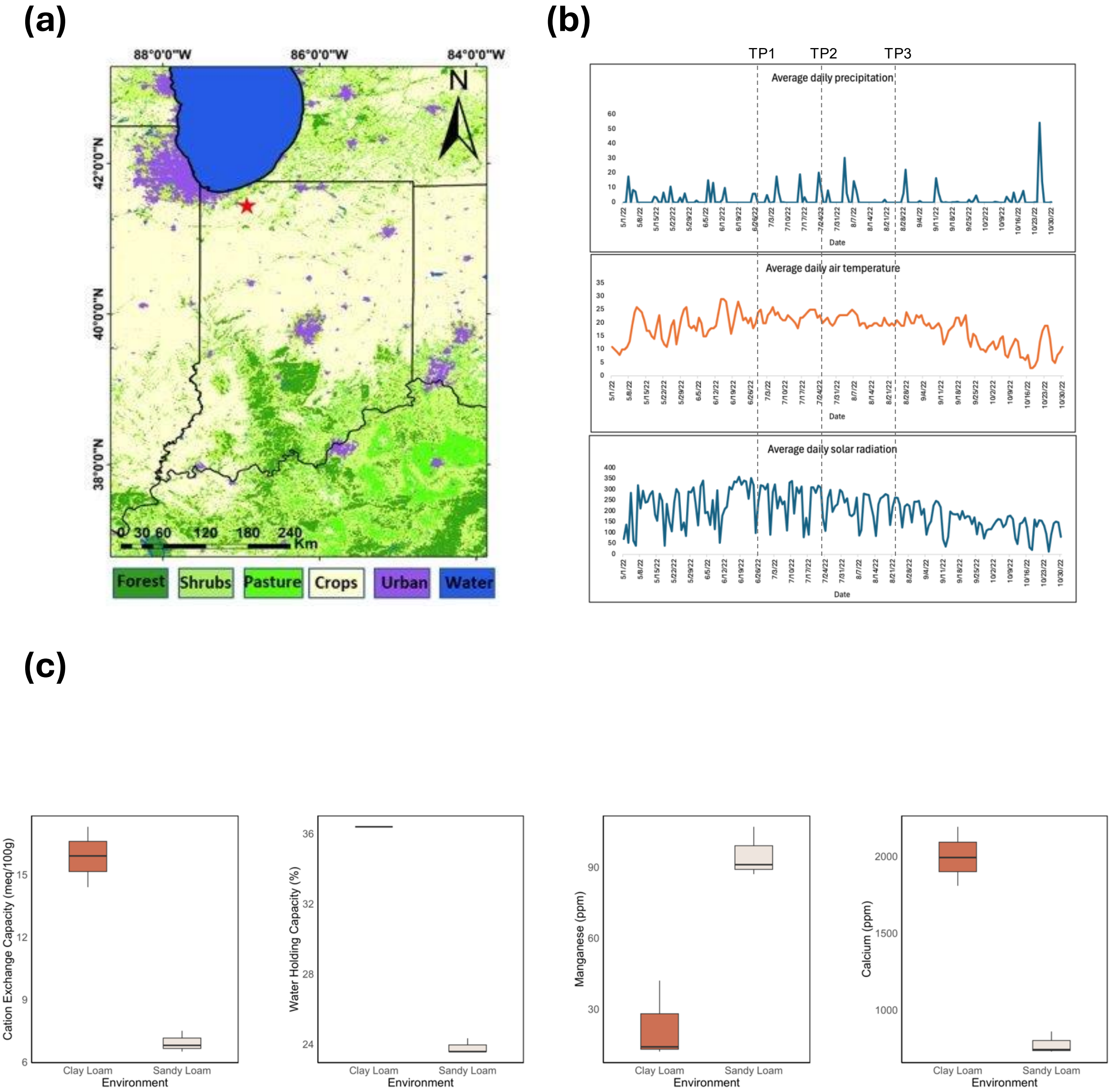


**Figure S1. Experimental site profile.** (a) Location of the experimental fields, highlighted by a red star. (b) Overview of weather data during the experimental season, showing average daily precipitation, temperature, and solar radiation. Dotted lines indicate three sampling timepoints. (c) Soil properties that differed between the two experimental sites.

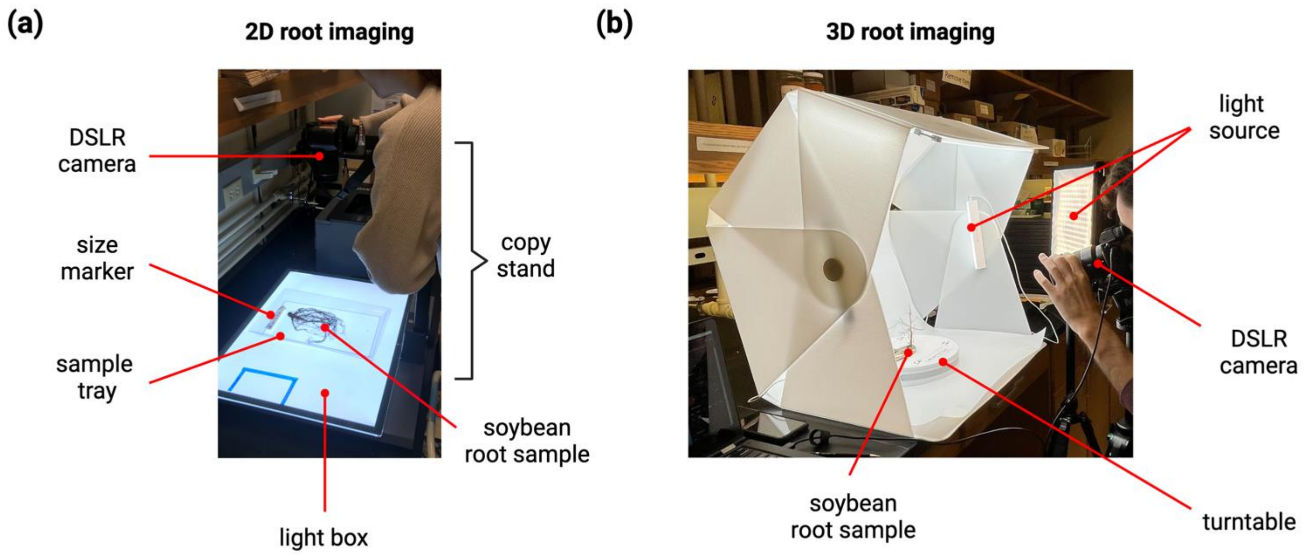


**Figure S2. Setup of root-imaging systems used in this study.** (a) 2D imaging system utilizing a copystand and lightbox. (b) 3D imaging system comprising of a turntable and light sources. Both systems utilized DSLR cameras that were operated manually on fixed platforms.

Extracting root traits from 2D images with ImageJ

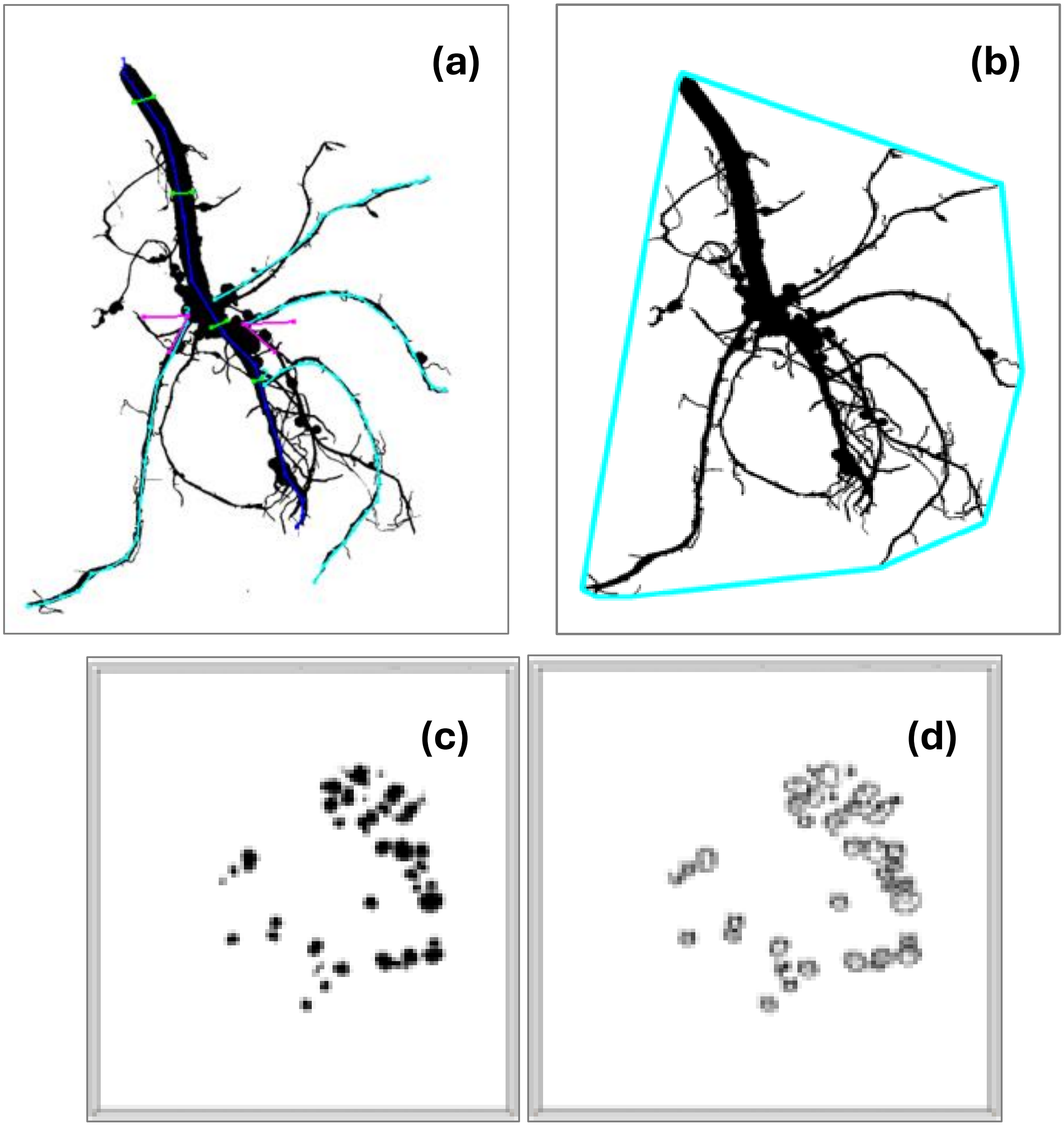


**Figure S3. Extracting root traits from 2D images with ImageJ.** Image analysis examples shown here are taken from the Breeder Line genotype, G17 showing (a) its binarized image, where tap root diameter (green), lateral branch length (cyan), tap root length (blue), and average root angles (magenta) were sampled; (b) root spread as represented by the area of the convex hull (cyan) containing the root system; (c) automated detection of dissected root nodules and (d) segmented images of detached nodules

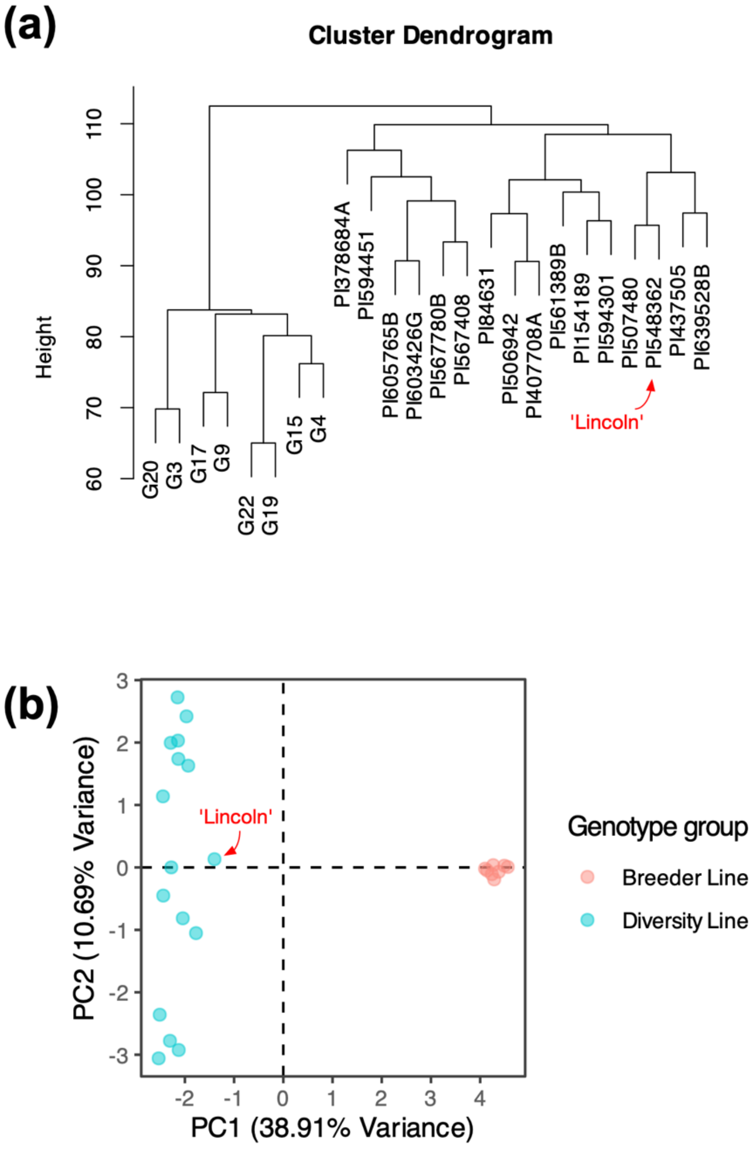


**Figure S4. Genetic structure of the soybean genotypes.** (a) Hierarchical clustering of soybean genotypes based on genetic distance. Genotypes are grouped into Breeder Line (Varieties' names starting from 'G' ) and Diversity Line categories (Varieties' names starting from 'PI'). (b) Principal Component Analysis (PCA) showing genetic variation across genotypes.

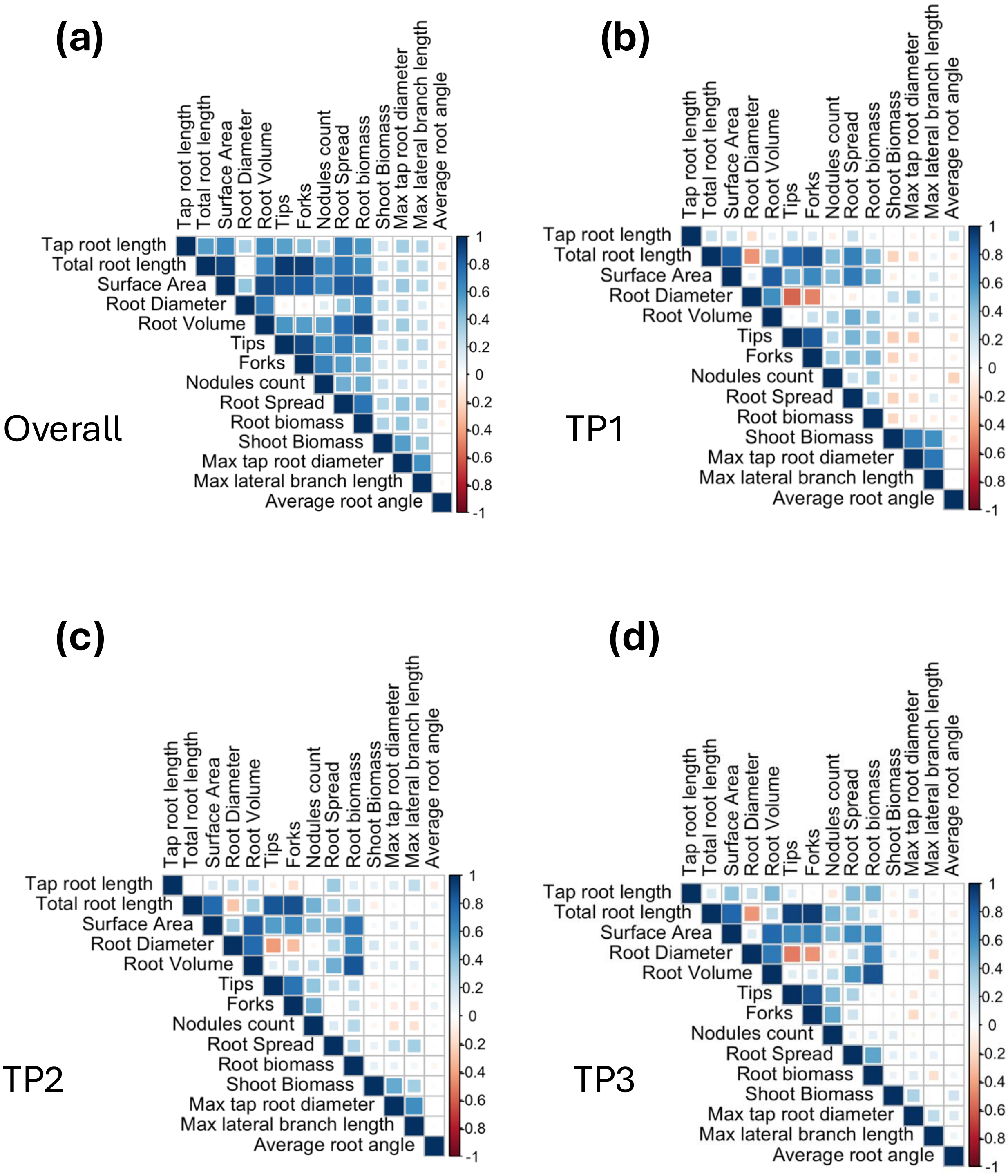


**Figure S5. Correlation of root traits and shoot biomass.** (a) Overall correlation heatmaps for root traits and shoot biomass. Correlation heatmap matrix from timepoints one (b), two (c), and three (d). The color reflects 'r' values and the size of the square indicates the p-value. Larger squares indicate lower p-values.

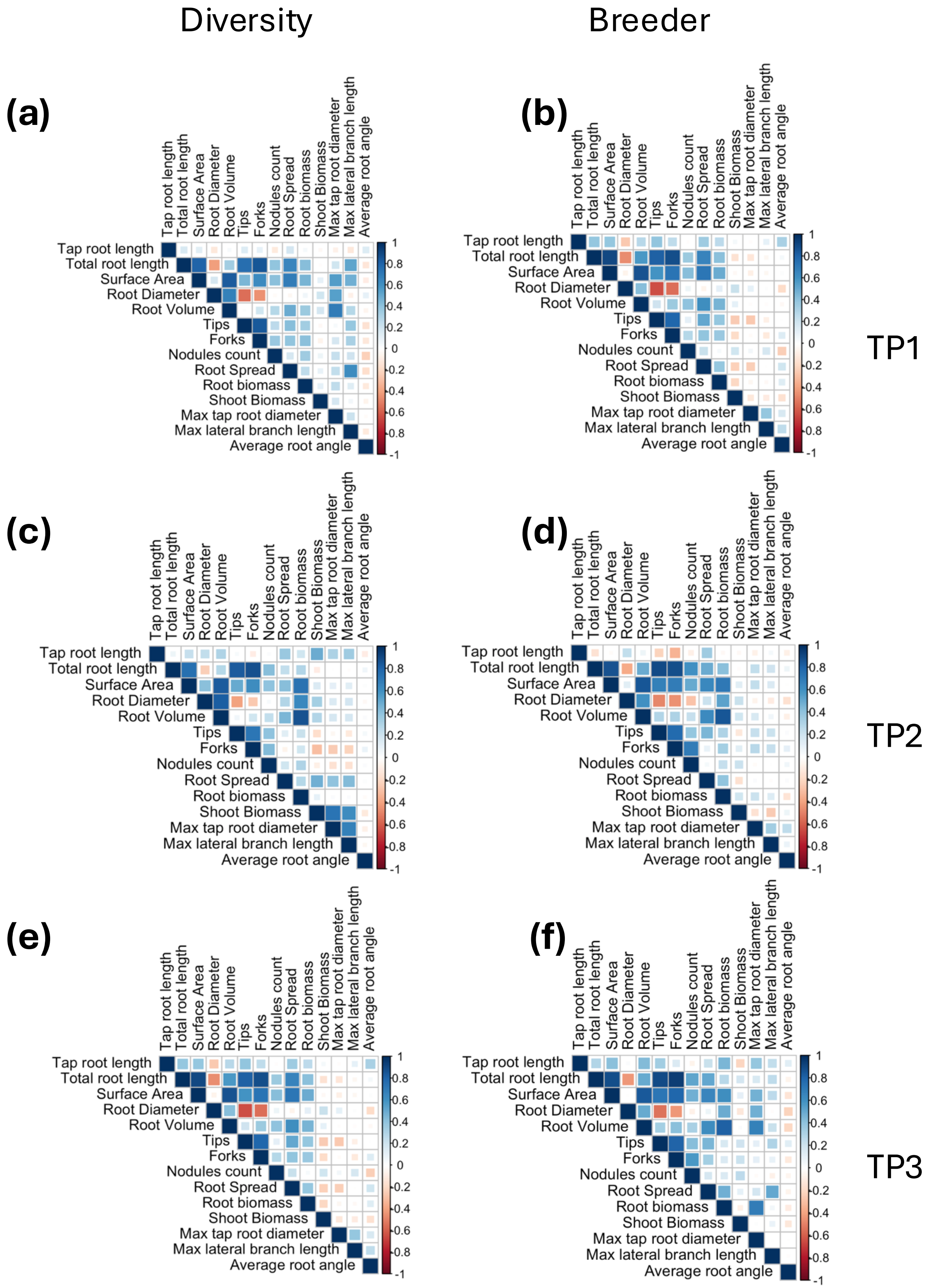


**Figure S6. Trait correlations across different combinations of timepoint and genotype group.** Correlation heatmaps plots shown for (a) TP1 diversity panel, (b) TP1 breeder panel, (c) TP2 diversity panel, (d) TP2 breeder panel, (e) TP3 diversity panel, and (f) TP3 breeder panel.

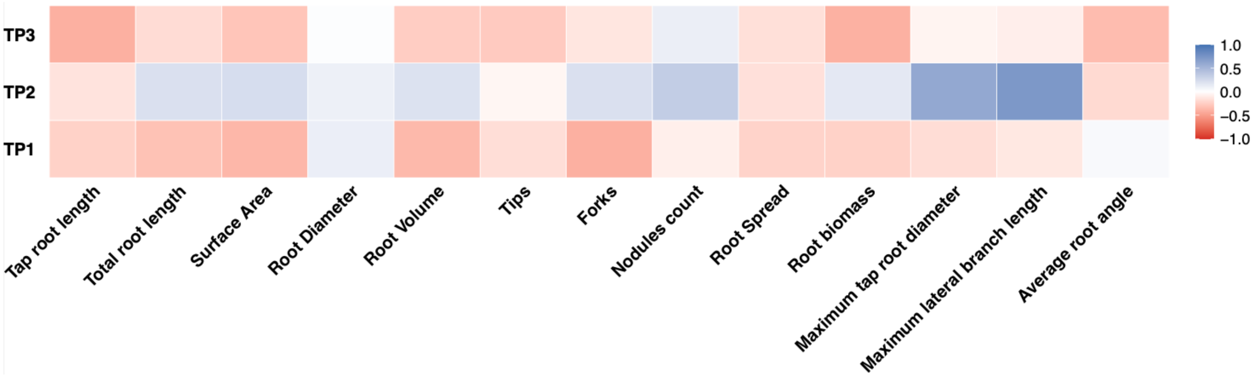


**Figure S7. Correlation between relative changes in shoot biomass and relative changes in root traits across environments at TP1 through TP3.** Heatmap visualization shows the correlation strengths over the three sampling timepoints, with the color scale indicating the strength of the correlation.

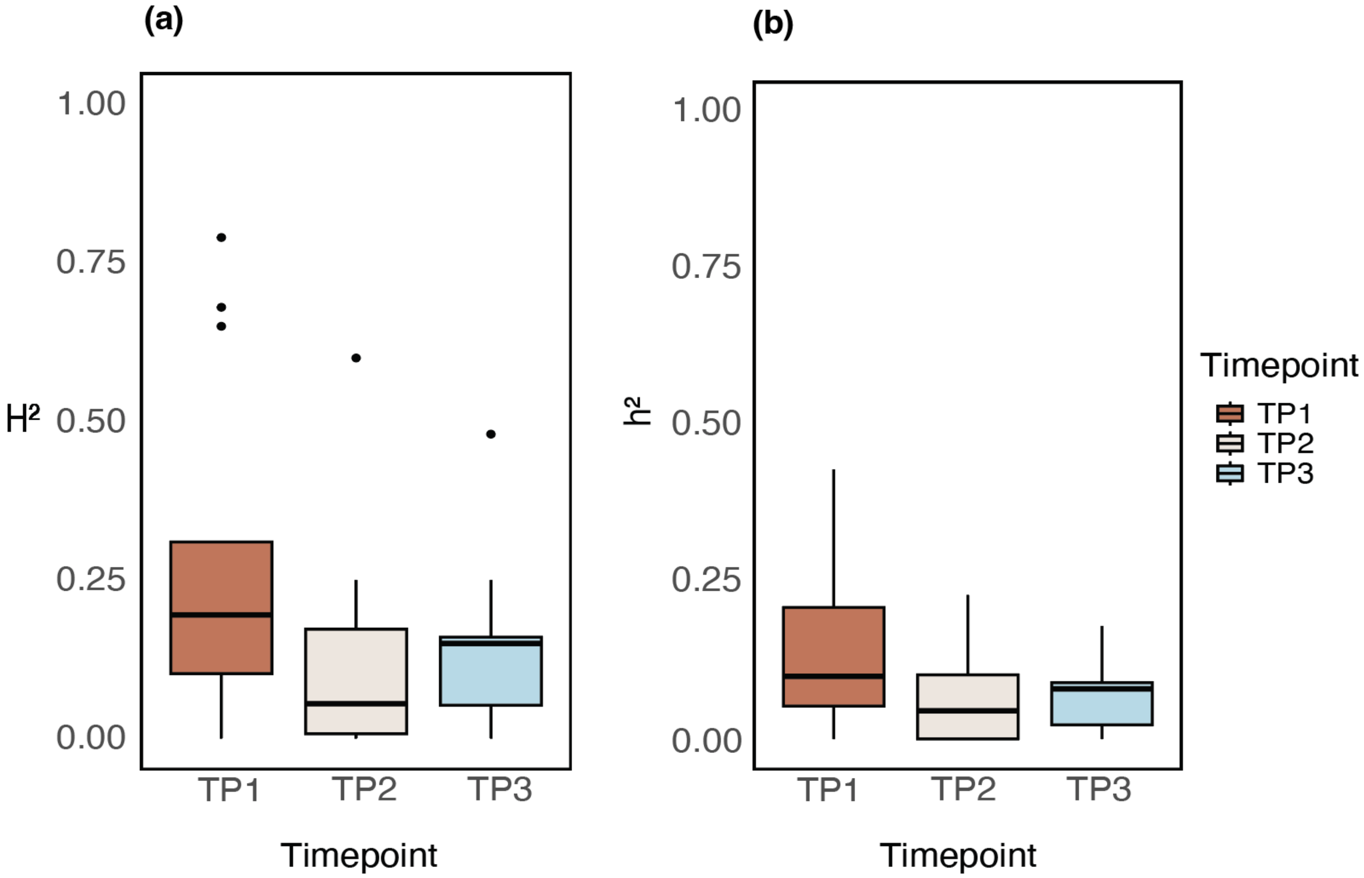


**Figure S8. Heritability of phenotypes of all traits across timepoints.** Boxplots show distributions of (a) broad-sense heritability ($H^2$) and (b) Narrow-sense heritability ($h^2$) for all traits (root and shoot).

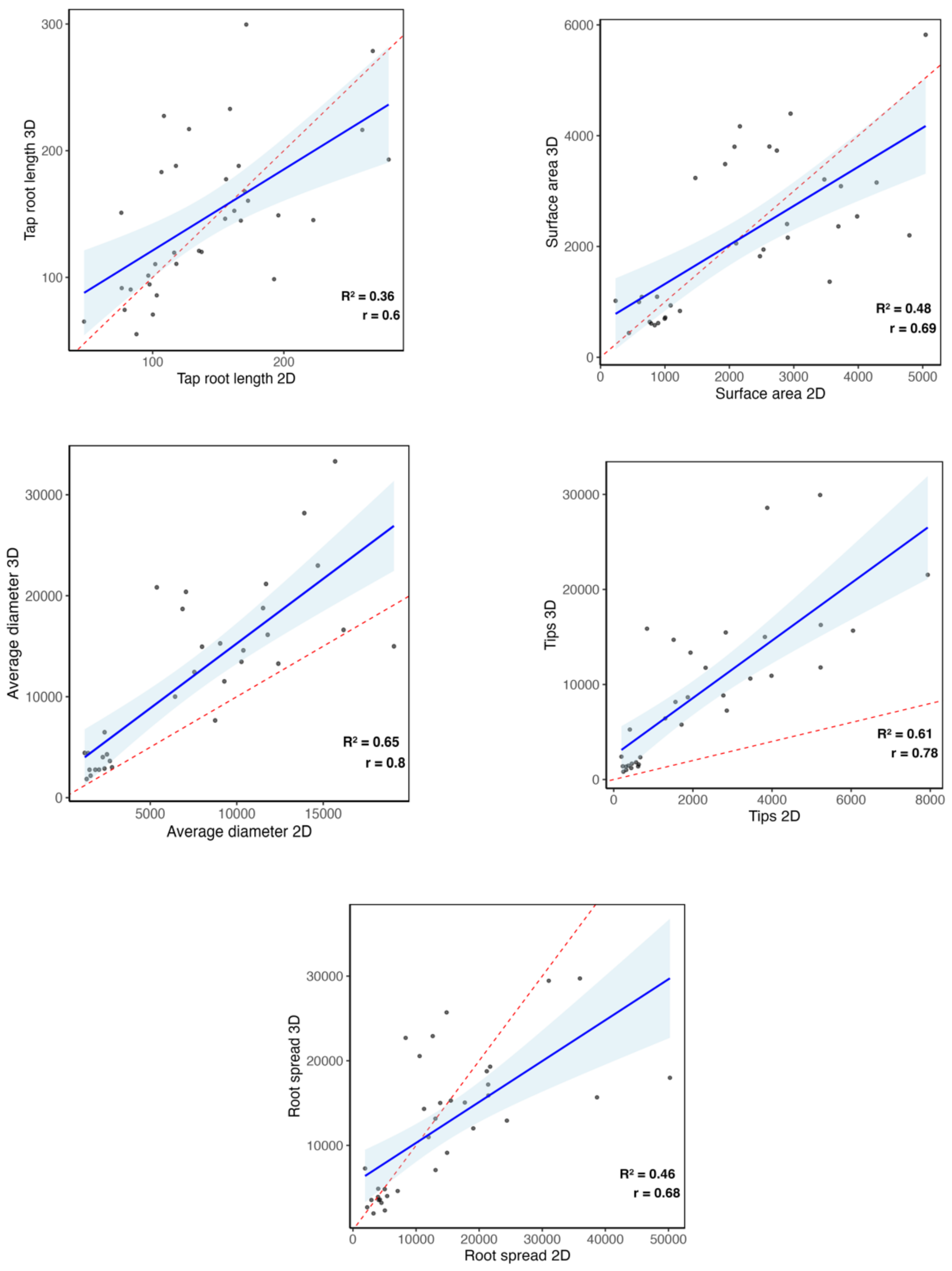


**Figure S9. Comparision of 2D versus 3D traits.** Subplots show comparisons of 2D (x-axis) and 3D (y-axis) measurements for root traits that had a correlation coefficient (r) of 0.6 or higher. The blue line shows the linear fit, and the red dashed line represents the 1:1 line. Traits include tap root length, surface area, average diameter, root spread, and number of tips.

Clay loam TP1

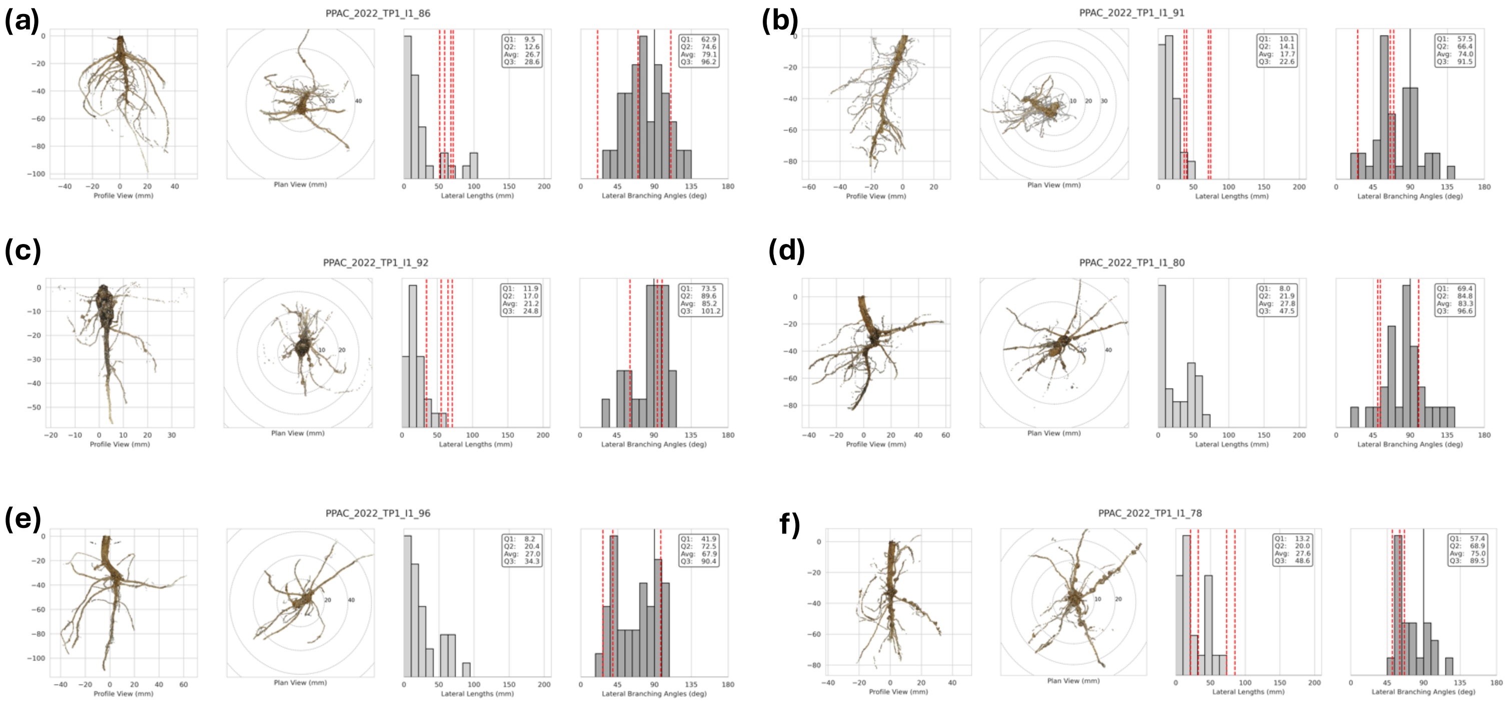


**Figure S10: Lateral root analysis from 3D reconstruction for samples from the clay loam environment in timepoint one.** In each panel, profile and plan views of root systems are shown along with histograms of lateral root lengths and branching angles. Dotted lines show values that were measured manually from 2D image analysis while the distributions represent the values derived automatically from 3D image analysis.

Sandy loam TP1

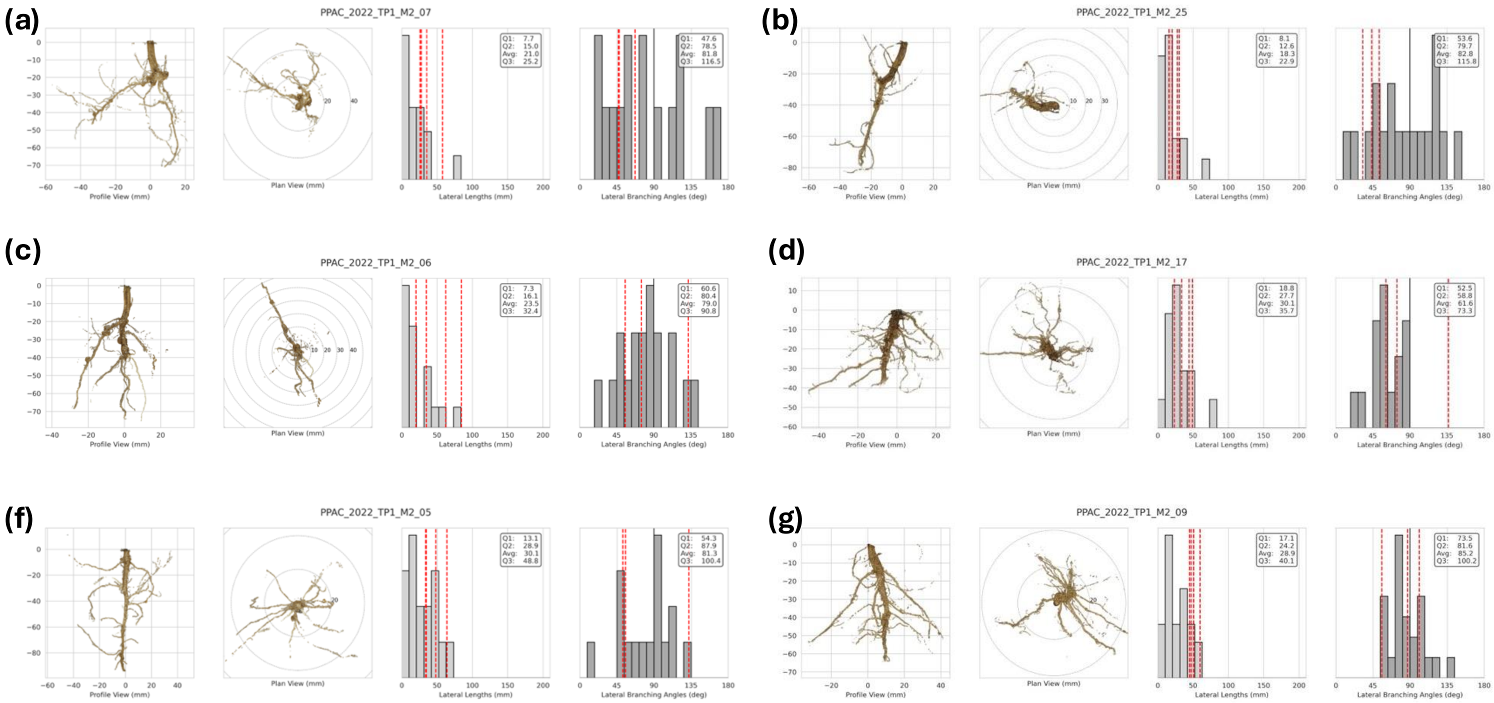


**Figure S11: Lateral root analysis from 3D reconstruction for samples from the sandy loam environment in timepoint one.** In each panel, profile and plan views of root systems are shown along with histograms of lateral root lengths and branching angles. Dotted lines show values that were measured manually from 2D image analysis while the distributions represent the values derived automatically from 3D image analysis.

Clay loam TP2

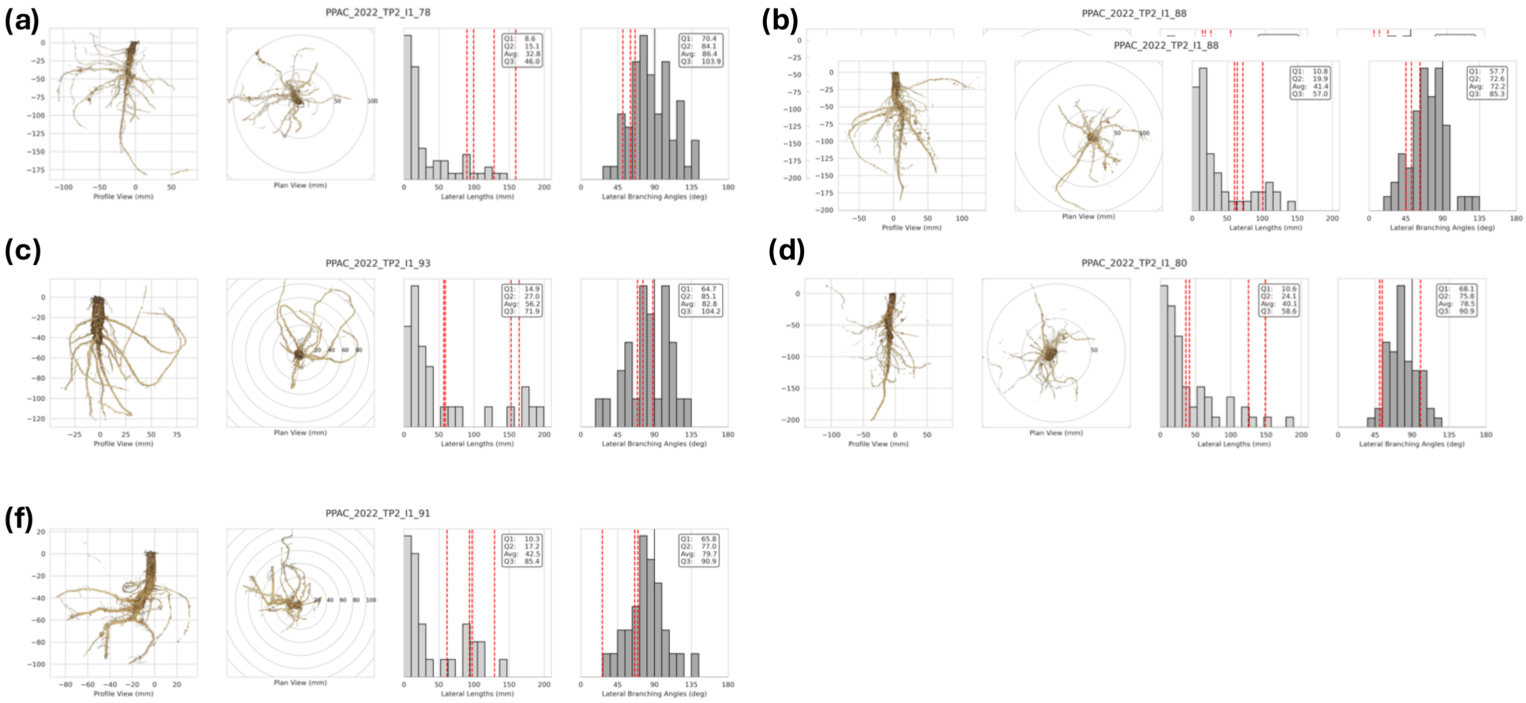


**Figure S12: Lateral root analysis from 3D reconstruction for samples from the clay loam environment in timepoint two.** In each panel, profile and plan views of root systems are shown along with histograms of lateral root lengths and branching angles. Dotted lines show values that were measured manually from 2D image analysis while the distributions represent the values derived automatically from 3D image analysis.

Sandy loam TP2

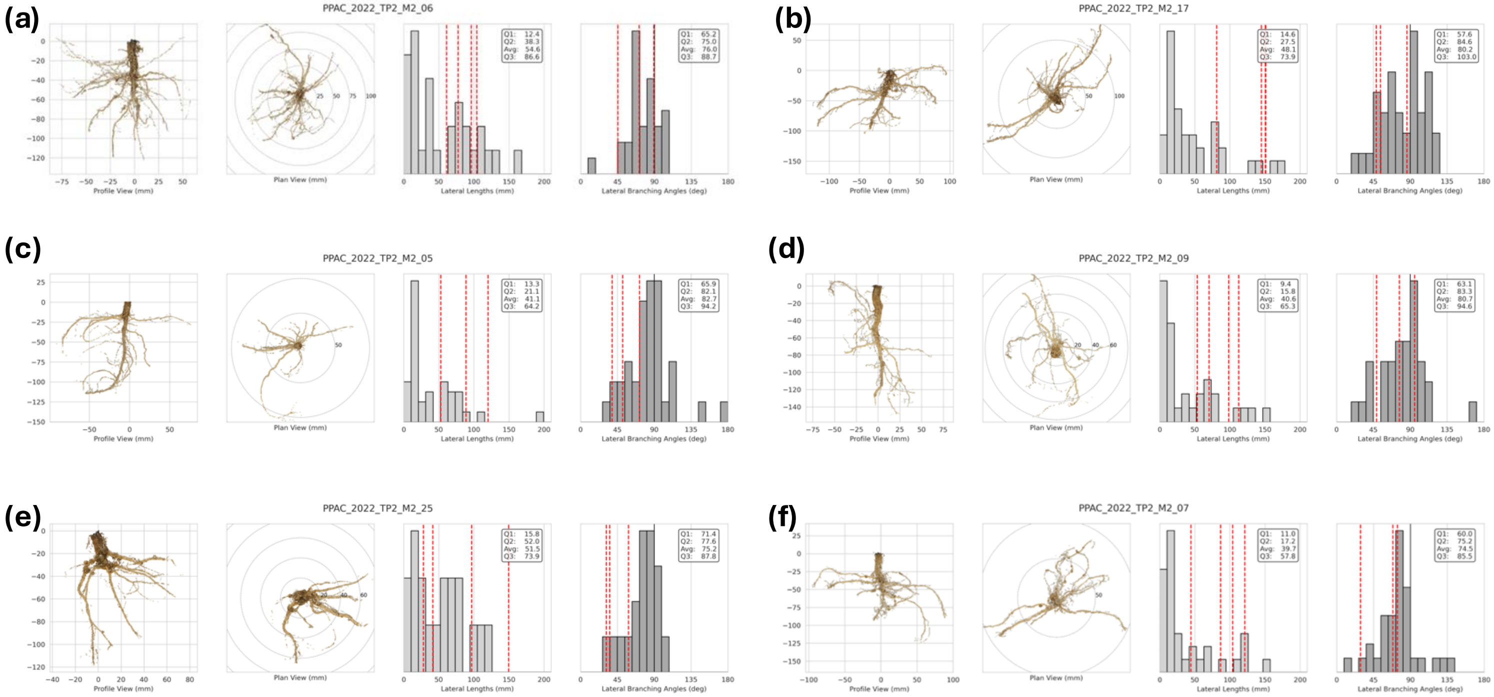


**Figure S13: Lateral root analysis from 3D reconstruction for samples from the sandy loam environment in timepoint two.** In each panel, profile and plan views of root systems are shown along with histograms of lateral root lengths and branching angles. Dotted lines show values that were measured manually from 2D image analysis while the distributions represent the values derived automatically from 3D image analysis.

Clay loam TP3

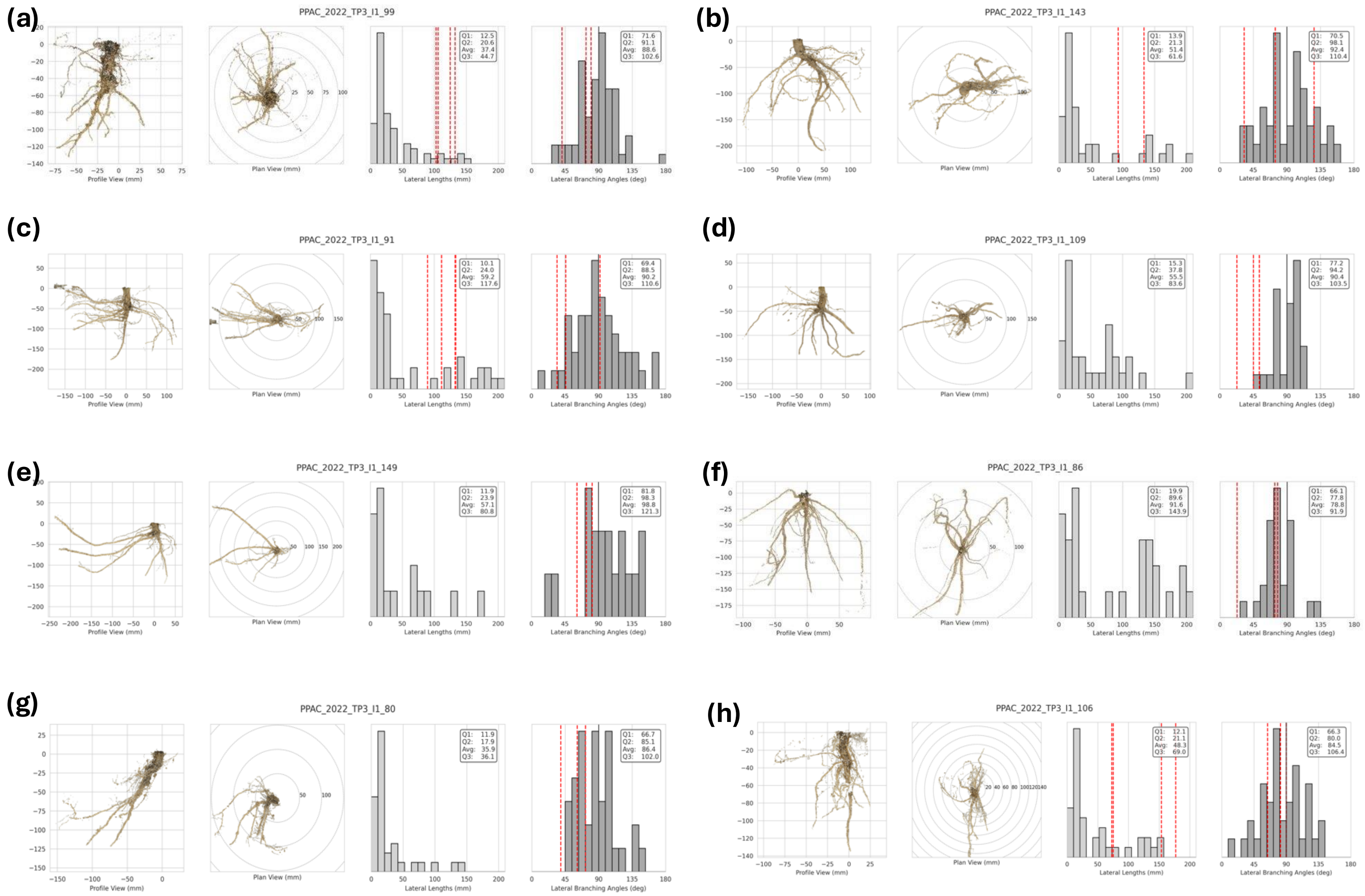


**Figure S14: Lateral root analysis from 3D reconstruction for samples from the clay loam environment in timepoint three.** In each panel, profile and plan views of root systems are shown along with histograms of lateral root lengths and branching angles. Dotted lines show values that were measured manually from 2D image analysis while the distributions represent the values derived automatically from 3D image analysis.

Sandy loam TP3

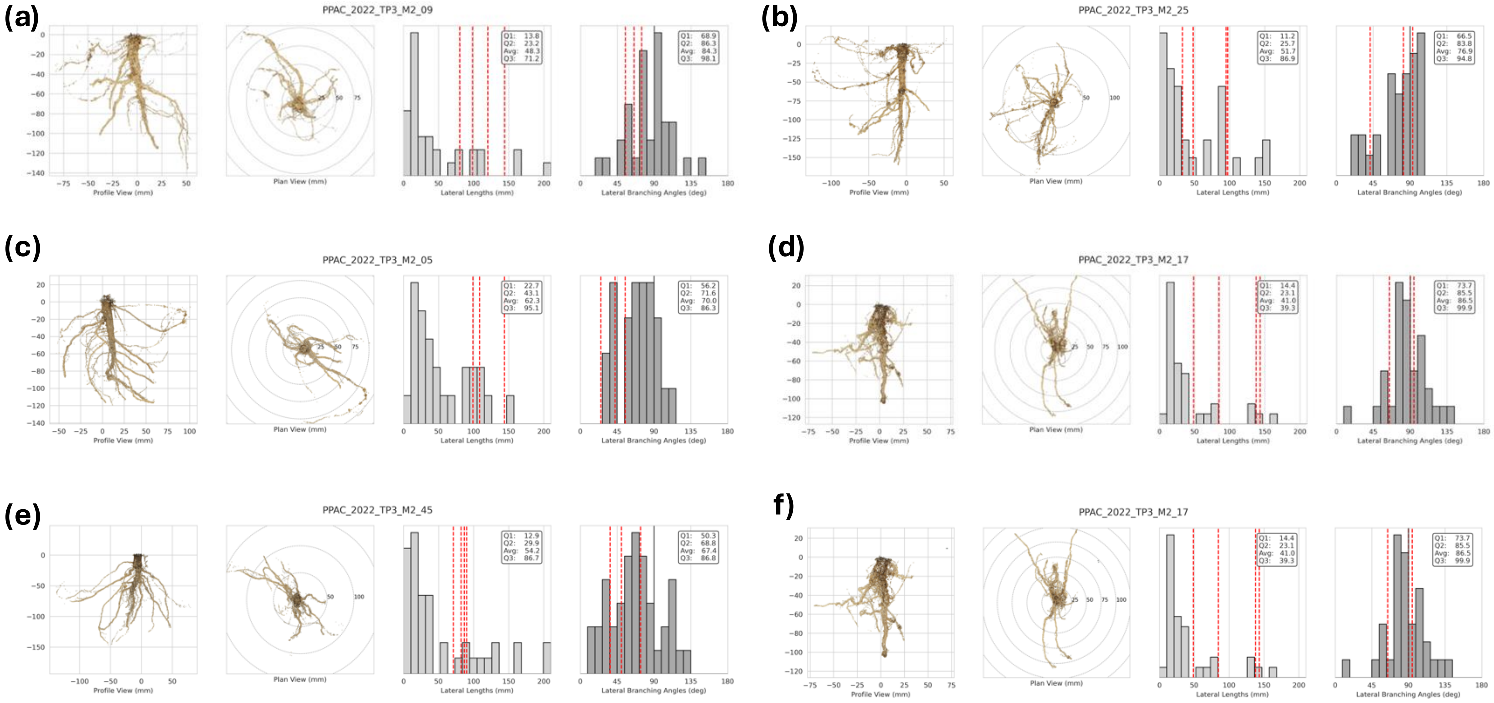


**Figure S15: Lateral root analysis from 3D reconstruction for samples from the sandy loam environment in timepoint three.** In each panel, profile and plan views of root systems are shown along with histograms of lateral root lengths and branching angles. Dotted lines show values that were measured manually from 2D image analysis while the distributions represent the values derived automatically from 3D image analysis.
